# Bottom-up engineering of EV-mimetic vesicles reveals selective protein loading and modulates cellular uptake and biodistribution

**DOI:** 10.64898/2026.10.05.756895

**Authors:** Yunxi Chen, Yousra Benslimane, Noélie Douanne, Kyle Dickinson, Thupten Tsering, Masoud Ghasemi, Laura Kienzle, Elisabet Frutos-Grilo, David Juncker, Anne-Laure Larroque, Luis Serrano Pubul, Vahé Nerguizian, Julia V. Burnier

**Affiliations:** Cancer Research Program, Research Institute of the McGill University Health Centre, Montreal, QC, Canada; Department of Pathology, McGill University, Montreal, QC, Canada; Department of Biomedical Engineering and Victor Philippe Dahdaeh Institute of Genomic Medicine, McGill University, Montreal, QC, Canada; Centre for Genomic Regulation, Barcelona, Spain; Proteomics and Molecular Analysis Platform, Research Institute of the McGill University Health Center, Montreal, QC, Canada; Universitat Pompeu Fabra (UPF), Barcelona, Spain; ICREA, Pg. Lluis Companys 23, 08010 Barcelona, Spain; École de technologie supérieure ÉTS, Montreal, QC, Canada; Gerald Bronfman Department of Oncology, McGill University, Montreal, QC, Canada

**Keywords:** extracellular vesicles, liposomes, biomimetics, delivery, nanomaterials

## Abstract

Biological systems such as extracellular vesicles (EVs) are highly sophisticated and functionally diverse. Despite growing recognition of EVs’ roles in health and disease through mediating intercellular communication, their inherent heterogeneity and challenges in purity and yield hinder their mechanistic research and clinical translation. To address this gap, we sought to translate essential EV properties into a controllable and simplified platform by engineering biomimetic synthetic vesicles (SVs) incorporating defined proteins and lipids. Using microfluidics, we generated SVs loaded with bulk membrane and cytosolic proteins from cancer cells, as well as individual organotropism-associated integrins. Bulk membrane protein incorporation into SVs reduced surface charge, modified lipid profiles, and enhanced cellular uptake, while integrin incorporation altered biodistribution. To investigate protein loading, we profiled input materials and SVs by mass spectrometry, identifying features linked to high incorporation, such as low molecular weight, positive charge, high isoelectric point, and predicted stability, globularity, lipidation, and amphipathic helices. This work presents a synthetic platform for dissecting the roles of EV-associated features and cargo, while providing insights into protein properties relevant to the optimization of delivery systems.

## Introduction

Extracellular vesicles (EVs) are lipid bilayer-enclosed vesicles naturally emitted by cells^1^, functioning as key mediators of intercellular communication through the transfer of diverse cargo, including proteins, lipids, and nucleic acids^2^. As EV research expands, an increasing diversity of EV subtypes and cargo profiles across physiological and pathological states continues to be uncovered, underscoring a dynamic and context-dependent molecular complexity^3^. For instance, in cancer, EVs play a pivotal role in shaping the tumor microenvironment and mediating organ-specific metastasis by establishing a premetastatic niche^4^. Our work and others have characterized cancer cell-derived EVs and identified candidate proteins, such as integrins, implicated in the metastatic process^5–7^.

With increasing understanding of their biological functions, EVs have attracted substantial interest as therapeutic agents and delivery systems^8^. Although numerous EV-based therapies have entered clinical trials over the past decade^9^, few have successfully reached the market. This limited translational success is primarily due to persistent challenges, which include identifying optimal biological sources and EV subtypes with desired functional properties^10^, scaling up EV production and purification to meet clinical-grade standards^11^, and delineating off-target effects caused by the inherent EV cargo^12^.

In contrast, synthetic liposomes, primarily developed as drug carriers, possess advantages in scalability and tunable composition^13,14^. Unlike EV biogenesis, liposome synthesis benefits from bottom-up fabrication, allowing for fine control over materials, size, and surface modifications^15^. However, the lack of biomolecules in liposomes limits their ability to recapitulate natural intercellular communication processes.

To overcome the challenges in EV research and biomedical applications, we propose a paradigm shift: the rational redesign of “EVs” by assembling only the biologically essential components into liposomes—yielding simplified, customizable, EV-like lipid bilayered synthetic vesicles (SVs) with defined composition. Several studies have explored strategies to integrate biological functionality with synthetic design. For example, semi-synthetic vesicles derived from bacterial outer membrane vesicles, with cytosolic content removed, have been explored to develop cancer vaccines with improved safety^16^. In another approach, liposomes functionalized with leukocyte membrane proteins have demonstrated enhanced circulation and homing to inflamed tissues^17–19^.

In the application of SVs in EV research, most studies on “EV-mimetics” have employed top-down approaches, such as physically disrupting cells into nanovesicles^20,21^. Despite increased yield, these methods retain biological complexity and continue to be challenged by high heterogeneity. To address these limitations, we generate SVs loaded with selected composition in a bottom-up manner using microfluidics. We generated and characterized diverse types of SVs loaded with either bulk membrane or cytosolic proteins from cancer cells, as well as SVs incorporating individual recombinant proteins, to demonstrate the modularity and precision of our approach. We systematically studied the physicochemical properties, cellular uptake, and biodistribution of a range of protein-functionalized SVs and their relevance to native EVs. To gain insight into the composition and assembly efficiency of the biomaterials (proteins and lipids), we performed comprehensive proteomic and lipidomic analyses and identified molecular features that influence protein-loading efficiency. While we mainly used cancer cell-derived materials, aimed at applying SVs as a model to advance research on cancer biology, we also extended to fabricating SVs with non-cancer materials and curcumin for demonstrating their utility in drug delivery.

Together, this work aims to establish a rational design framework for bottom-up construction of EV surrogates, offering a versatile platform to dissect EV biology and advance precision delivery systems.

## Results

### Physicochemical Characterization of Protein-Loaded SVs and Their EV-Mimicry

We previously engineered liposomes mimicking the physicochemical properties of cancer-derived EVs, synthesizing SVs with predicted size and charge to systematically study their influence on cellular uptake^22^. Building on this approach, we now functionalize SVs with biological cargo. As a proof-of-concept, whole cell lysate (WCL) extracted from MP41, a uveal melanoma cell line with known liver metastatic potential, was loaded into liposomes (WCL-SVs) (**Supplementary Data 1**). Both Empty-SVs (liposomes devoid of proteins) and WCL-SVs exhibited size distributions resembling small EVs (sEVs), predominantly below 200 nm (**Supplementary Data 1a**). Cryogenic transmission electron microscopy (Cryo-TEM) confirmed lipid bilayer structure, verifying liposomal identity (**Supplementary Data 1b**). Selected proteins, including ITG AV, actin, and canonical sEV-associated markers (TSG101, CD81, and HSP90), were successfully detected in WCL-SVs (**Supplementary Data 1c**). Interestingly, WCL-SVs demonstrated significantly enhanced cellular uptake compared to Empty-SVs (**Supplementary Data 1d-g**), indicating a crucial role of proteins in nanoparticle-cell interactions. These structural, molecular, and functional validations underscore the potential of protein-loaded liposomes to mimic EVs, inspiring further tuning of SVs’ composition to modulate their functions and investigate their EV-mimicry. Specifically, we hypothesized that membrane-associated proteins within the WCL fraction predominantly mediated cellular uptake.

To test this hypothesis, cell membrane protein-enriched (MemRaw) and cytosolic protein-enriched (CytoRaw) fractions were separately isolated from MP41 cells using a commercial Mem-PER™ Plus kit and loaded into SVs during microfluidic synthesis. Four types of SVs were generated: Empty-SVs, Mem-SVs (MemRaw loaded), Cyto-SVs (CytoRaw loaded), and Complete-SVs (containing both MemRaw and CytoRaw). sEVs were extracted from the same cell line MP41. The sEVs and SVs were characterized for their physical and functional properties, including size, charge, morphology, proteome and lipidome, cellular uptake, and biodistribution (**Fig. 1a**).

**Figure 1.**
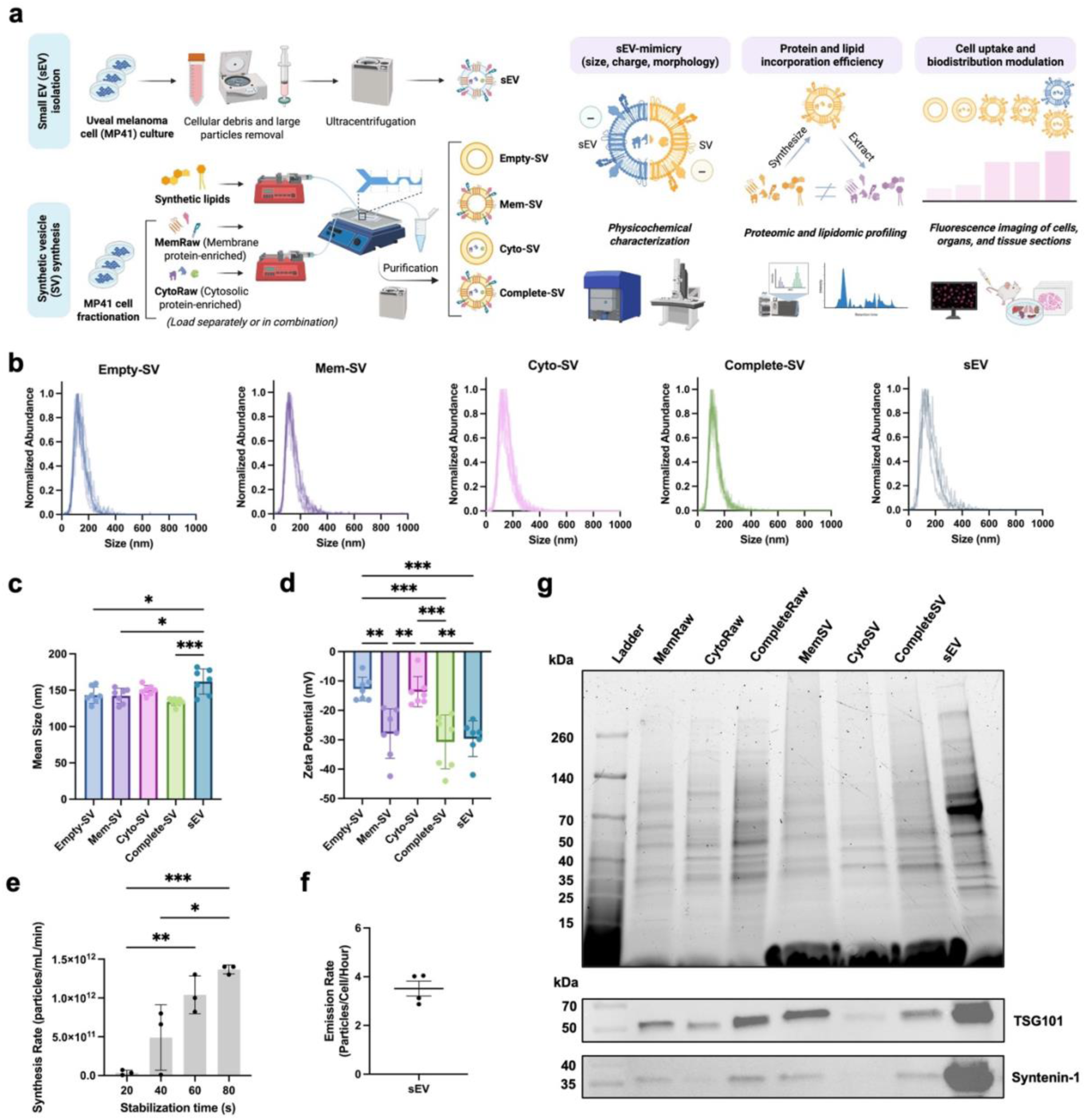
Study design and physical characterization of small EVs (sEVs) and synthetic vesicles (SVs). **a**, Methodologies of sEV isolation, SV synthesis, and characterization. sEVs and SVs were characterized for size distribution (the particle concentration was normalized to the maximum concentration in each distribution to overlay distributions from multiple independent repetitions to the same 0 to 1 scale) (**b**), mean size (**c**), zeta potential (**d**), synthesis rate of SVs (**e**), and emission rate of sEVs (**f**). **g**, SDS-PAGE gel of total protein bands in raw proteins, SVs and sEVs, and western blots for TSG101 and Syntenin-1. **b-f**, Biological replicates. One-way ANOVA followed by Tukey’s multiple comparison test. Mean ± SD. *, **, and *** represent *P* < 0.05, 0.01, and 0.001, respectively.

All SVs and sEVs exhibited a mean size in the range of 100 – 200 nm, measured by ZetaView (**Fig. 1b-c**). However, it is noted that TEM analysis indicated that SVs generally exhibited larger apparent diameters, approximately 100–300 nm, whereas sEVs contained a higher proportion of smaller particles, many within the 20–100 nm range (**Supplementary Data 1h**). These technique-specific size differences could be because TEM captures vesicle morphology under dehydrated and vacuum conditions that cause shrinkage of EVs, as previously reported^23^, whereas nanoparticle tracking analysis (NTA), like ZetaView, measures hydrated particles in suspension. Natural EVs and SVs may respond differently to these measurement environments, due to differences in lipid composition that influence membrane rigidity and luminal content, which could contribute to the observed size differences. In addition, NTA has limited sensitivity for vesicles below approximately 70 nm, which could shift the apparent mean size of the sEV population upward^24^.

Incorporation of MemRaw significantly made the zeta potential (surface charge) of SVs more negative, while incorporation of CytoRaw did not have this effect (**Fig. 1d**). This electrostatic shift aligns with the natural surface properties of EVs and supports the surface-mimicry of Mem-SVs. At a stable synthesis rate, SVs were generated at concentrations exceeding 1E12 particles/mL/min (**Fig. 1e**), while the mean emission rate of isolated sEVs was approximately 5 particles/cell/hour (**Fig. 1f**). Protein bands were detected in Mem-SVs, Cyto-SVs, and Complete-SVs, validating protein incorporation (**Fig. 1g**).

Exosomal markers (TSG101, syntenin-1) were validated in the sEVs, providing benchmarks for evaluating sEV-mimetic cargoes of the SVs (**Fig. 1g**). Interestingly, these non-membrane EV marker proteins were more abundant in MemRaw and Mem-SVs than in CytoRaw and Cyto-SVs, potentially due to the solubilization of organelle membranes of TSG101 and syntenin-1-enriched organelles, such as endosomes, during membrane protein extraction. TSG101 and syntenin-1 are cytosolic adaptor proteins that associate with lipid membranes through protein– protein and lipid-mediated interactions. TSG101 binds hepatocyte growth factor-regulated tyrosine kinase substrate (HRS) on early endosomes^25^, while syntenin-1 is a cytosolic adaptor protein that is recruited to the plasma membrane through interactions with partner proteins and specific phosphoinositides^26^.

Collectively, these results demonstrate that SVs possess uniform and reproducible physicochemical characteristics and can be synthesized with protein cargoes in an efficient manner.

### Proteomic Profiling of SVs and Validation of Surface Protein Incorporation

Protein packaging into liposomes is known to be technically challenging. Elucidating the underlying mechanisms is essential for the rational design of synthetic EVs with defined biological functions. To bridge these gaps, we conducted mass spectrometry on proteins extracted from SVs (MemSVs, CytoSVs, and CompleteSVs), and their respective raw protein extracts prior to microfluidic synthesis (MemRaw, CytoRaw, and CompleteRaw). By correlating proteomic profiles with physicochemical properties, we identified key factors governing SV cargo loading (**Fig. 2a**).

**Figure 2.**
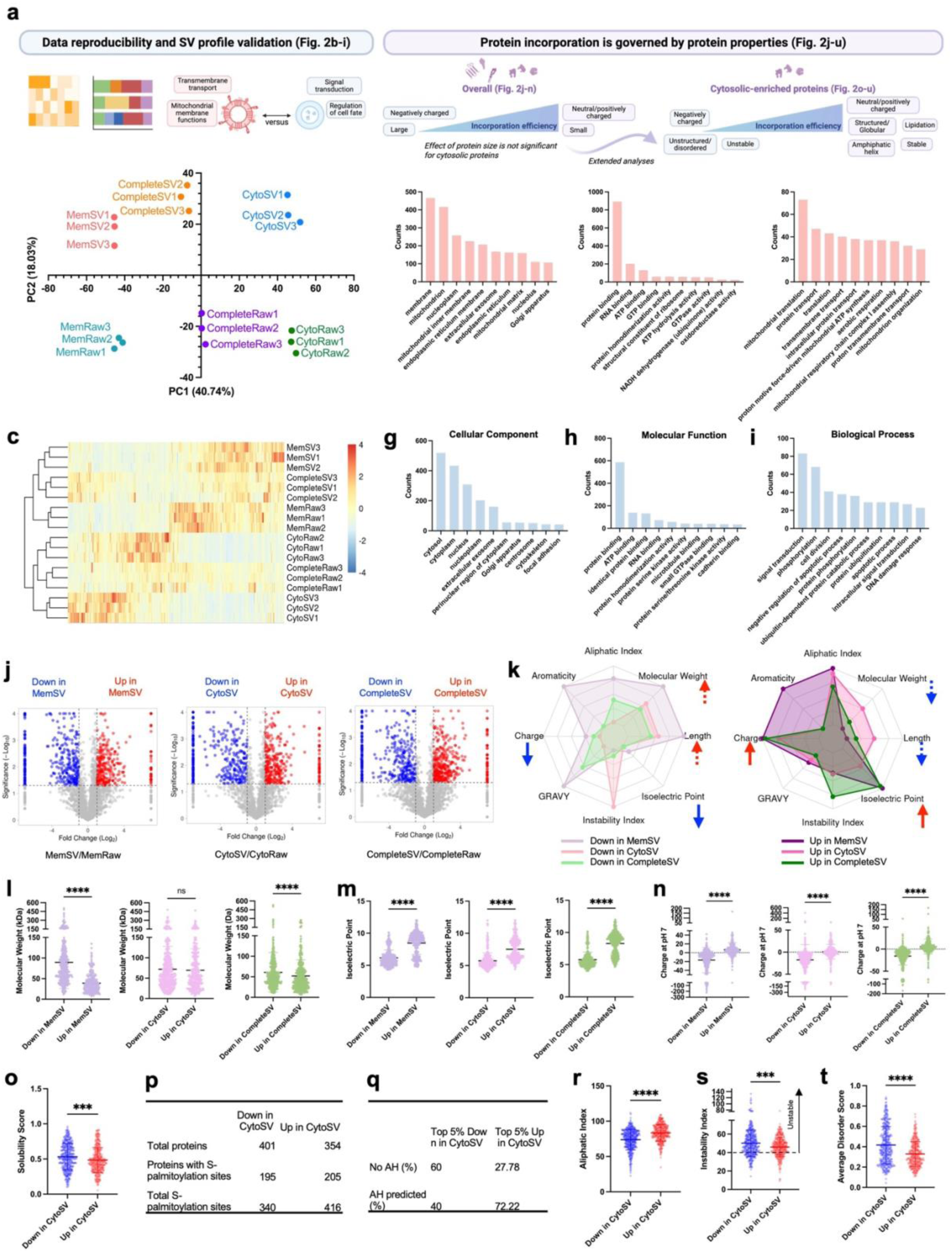
Proteomic profiling of SVs and their raw proteins. **a**, Schematic summarizing the results. Principal component analysis (**b**) and heatmap (**c**) show clustering profiles of biological triplicates. **d-i**, Gene ontology analysis. Top 10 terms of cellular components, molecular functions, and biological processes of unique proteins in MemSVs (**d-f**) and CytoSVs (**g-i**) were shown. **j**, Volcano plots compared SVs to their raw proteins. Proteins with *P* value < 0.05 and fold change < 0.5 (Down in SVs) and > 2 (Up in SVs) were considered significant. **k**, Radar plot summarizes median values on a scale of min to max of protein properties of significantly Down and Up-incorporated proteins for MemSV/CytoSV/CompleteSV. Comparisons of proteins in Down and Up in MemSV/CytoSV/CompleteSV for their protein molecular weight (**l**), isoelectric point (**m**), and net charge at pH7 (**n**). Comparisons of proteins in Down and Up in CytoSVs for solubility score (**o**), S-palmitoylation prediction (**p**), percentage of proteins with predicted amphipathic helix (AH) (**q**), aliphatic index (**r**), instability index (**s**), and average disorder score (**t**). **l-o**, **r-t**, Mann–Whitney U test. Mean ± SD. ns, ***, and **** represent *P* > 0.05, <0.001, and <0.0001, respectively.

Principal component analysis and heatmaps confirmed the clustering of biological triplicates, demonstrating high experimental reproducibility (**Figs. 2b-c**). Between 2486 and 3513 unique proteins were identified in each type of sample (**Supplementary Data 2a**). Despite the overlap between MemRaw, CytoRaw, and CompleteRaw, 1285 and 731 proteins were exclusively detected in MemRaw versus CytoRaw, respectively. 1188 and 702 proteins were exclusively detected in MemSV versus CytoSV, respectively. Venn diagrams of presence and absence revealed that SVs contain 85% to 88.2% of protein identities from their corresponding raw proteins (**Supplementary Data 2a**).

Proteins uniquely identified in MemSVs compared to CytoSVs were primarily associated with the plasma membrane, mitochondria, and nucleoplasm (**Figs. 2d-f**). These proteins displayed molecular functions in protein, RNA, ATP, and GTP binding and participated in biological processes such as mitochondrial translation, protein transport, transmembrane transport, and translation regulation. In contrast, proteins uniquely enriched in CytoSVs compared to MemSVs were predominantly associated with the cytosol, cytoplasm, nucleus, and nucleoplasm (**Figs. 2g-i**). These proteins exhibited functions related to protein, RNA, and ATP binding, as well as protein serine kinase activity. They were involved in biological processes such as signal transduction, phosphorylation, cell division, apoptotic regulation, and DNA damage responses, suggesting roles in intracellular signaling and stress response pathways. This compartment-specific functional enrichment demonstrates SVs’ ability to recapitulate and tailor native biological profiles—a key advantage over conventional EV studies limited by heterogeneity.

The expression patterns of selected proteins validate enrichment of individual membrane and cytosolic proteins (**Supplementary Data 2b**). Representative membrane-associated proteins, including cluster of differentiation (CD) markers (CD44, CD47, CD55, CD59, CD63, CD68, CD81), integrins, G-protein-coupled receptors (GPR107, GPR143, GPR161, GPR180), and Insulin-like growth factor 1 receptor (IGFR), were more enriched in MemRaw compared to CytoRaw. In contrast, cytosolic proteins such as GAPDH, beta-tubulin (TUBB), and S100A11 were predominantly enriched in CytoRaw. Interestingly, differential enrichment within the HSP90 family was observed: HSP90AA1 was enriched in CytoRaw, consistent with its cytosolic localization, while HSP90B1 was enriched in MemRaw, in line with its known residence in the endoplasmic reticulum lumen and occasional membrane association^27^. Enrichment of SDCBP (syntenin-1) and TSG101 in MemRaw and MemSVs, likely due to the co-extraction of organelle proteins during membrane solubilization, corroborates the western blot result (**Fig. 1g**). Overall, these results demonstrate the distinct profiles of MemRaw/SV versus CytoRaw/SVs.

Additionally, key markers of uveal melanoma, including MLANA, PMEL, and TYRP1, were identified, confirming the MP41 cell line as the protein source (**Supplementary Data 2b**). MP41 cells harbor the characteristic uveal melanoma driver mutation in the *GNA11* gene, which leads to constitutive activation of downstream oncogenic signaling pathways^28,29^. Consistent with this, GNA11 protein was detected in our analysis. Typical exosomal markers^12^ such as CD63, CD81, syntenin-1 (SDCBP), and TSG101 were also identified, supporting the biological relevance of SVs as cancer cell-derived, EV-mimetic vesicles.

The observed variation in protein incorporation efficiency through a heatmap prompted us to investigate its correlation with intrinsic protein properties. We subsequently identified significantly preferentially (Up) and poorly (Down) incorporated proteins (**Fig. 2j**) and analyzed their physicochemical properties. SV protein incorporation was found to be a multifactorial process influenced by diverse physicochemical attributes, as evidenced by the distinct profiles of proteins Up versus Down in SVs (**Figs. 2k-n, Supplementary Data 2c-d**).

Protein size and electrostatic interactions appeared as key drivers of SV protein incorporation. Proteins Up in MemSVs displayed significantly smaller mean molecular weights (39.10 ± 22.33 kDa) compared to proteins Down in MemSVs (89.55 ± 63.57 kDa, *P* < 0.0001) (**Fig. 2l**), suggesting that steric hindrance limits the incorporation of larger proteins. While the same trend exists for CytoSVs, it was not significant, underlying the intrinsic differences in the membrane and cytosol-derived proteins. Proteins Up in all SVs consistently exhibited significantly higher mean isoelectric points and higher net positive charges at pH7 (*P* < 0.0001 for all) (**Fig. 2m-n**). To account for the partial overlap of cytoplasmic organelle-associated proteins between MemRaw and CytoRaw fractions, we further annotated proteins for “transmembrane” or “cytosolic but non-transmembrane” using UniProt. The effect of these factors was reanalyzed and validated (**Supplementary Data 3-4**).

Incorporation of cytosolic proteins was found to be related to hydrophobicity, post-translational modifications, structural motifs, and stability. Given that cytosolic proteins are typically less dependent on lipid interactions for stability, their incorporation provides a more direct view of protein-intrinsic features affecting SV loading. It therefore prompted us to focus on the analysis of cytosolic proteins. Compared to proteins Down in CytoSVs, proteins Up in CytoSVs exhibited significantly higher GRAVY (grand average of hydropathicity of amino acids) score (*P* < 0.05) (**Supplementary Data 2c**), indicating stronger hydrophobic character. This correlated with lower predicted solubility scores (*P* < 0.001) (**Fig. 2o**), suggesting hydrophobicity facilitates spontaneous partitioning into lipid membranes during vesicle formation. Moreover, Up in CytoSVs proteins exhibited higher predicted S-palmitoylation rates (**Fig. 2p**). This lipid-anchoring modification could enhance membrane affinity through covalent attachment to palmitate molecules in the lipid bilayer^30^. 72.22% of the top 5% Up proteins showed strong predictions for amphipathic helices, compared to only 40% among the top 5% most Down proteins (**Fig. 2q**, **Supplementary Data 5–6**). This suggests a role for amphipathic helices in mediating membrane association, where the hydrophobic face of the helix inserts into the lipid bilayer while the hydrophilic face remains solvent-exposed^31^. Such structural features enable stable peripheral membrane association without requiring full transmembrane integration, supporting a model of selective protein-lipid interaction that facilitates cytosolic protein incorporation into SVs.

Protein stability represents another key factor for cytosolic protein incorporation. Proteins Up in all SV types consistently exhibited significantly higher aliphatic indices, a parameter associated with increased thermostability^32^ (**Supplementary Data 2d**). This effect was especially pronounced in CytoSVs (*P* < 0.0001) (**Fig. 2r**). Additionally, proteins enriched in CytoSVs had significantly lower mean instability indices (*P* < 0.001) (**Fig. 2s**) and a lower disorder score (**Fig. 2t**), indicating a preference toward incorporating structurally stable proteins in CytoSVs. These stability trends may favor retention of functional cargos in SVs. Collectively, these findings highlight the significant influence of physicochemical factors on protein incorporation efficiency. These findings could provide actionable criteria for optimizing protein loading in therapeutic SVs.

In terms of EV-mimetic assessment, comparison with the MP41 sEV label-free mass spectrometry dataset from Tsering et al.^6^ showed that 73.37% of proteins identified in MP41 sEVs were also detected in CompleteSVs (**Supplementary Data 2e**). In addition, 2350 proteins were uniquely detected in CompleteSVs, likely reflecting the broader total cellular protein input used for SV generation, together with differences in sample preparation and technical variation. Notably, the overlapping protein set was significantly enriched for exosome- and extracellular vesicle-related cellular component Gene Ontology terms (**Supplementary Data 2f**), supporting partial proteomic mimicry of natural sEVs by CompleteSVs.

To confirm the successful incorporation of surface proteins into SVs, multiple complementary techniques were employed. We specifically focused on detecting ITG AV because of the well-established role of integrins in mediating cancer organotropism^5^, as well as the relatively high abundance of ITG AV in the SVs compared to other integrins, which may facilitate detection (**Supplementary Data 7a**). Spectral counts (**Fig. 3a**) and western blot analysis (**Figs. 3b–c**) demonstrated effective incorporation of ITG AV into Mem-SVs and Complete-SVs, with lower detection in CytoSVs, as expected. Importantly, negative controls synthesized without lipids showed minimal protein detection, confirming efficient removal of unincorporated proteins and fidelity of SV-encapsulated protein detection (**Fig. 3b)**.

**Figure 3.**
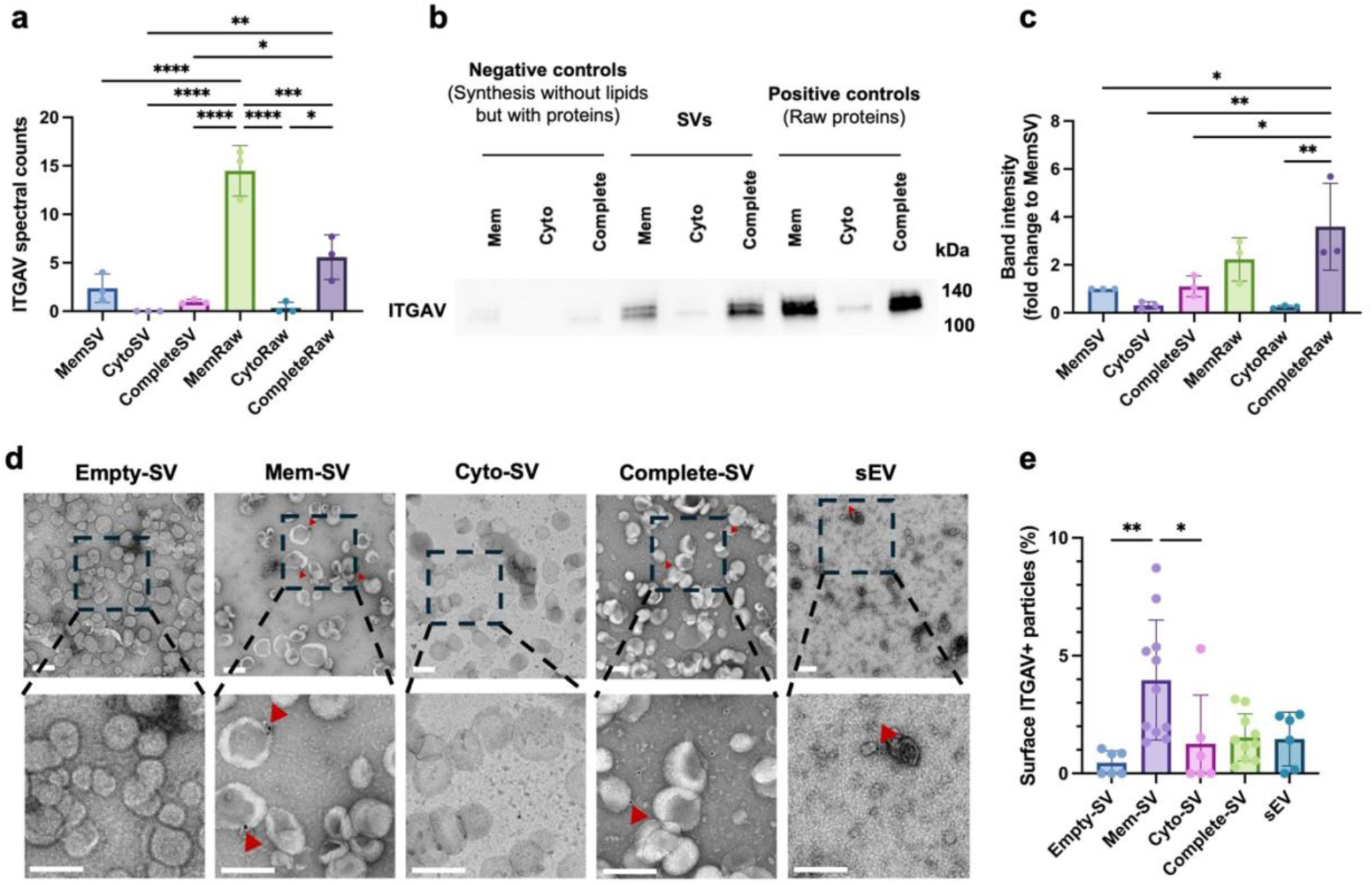
Validation of surface protein incorporation. **a**, Spectral counts of integrin αV (ITG AV) across SVs and raw proteins, measured at equal total input protein amount by mass spectrometry (n=3 syntheses). **b**, Western blot of ITG AV in proteins extracted from SVs, raw proteins as positive controls, and synthesis with proteins but no lipids as negative controls. **c**, Quantification of relative band intensity across SVs and raw proteins, shown as fold change to MemSV (n=3 independent experiments). **d**, Immunogold labeling of SVs and sEVs against ITG AV. Scale bars = 200 nm. **e**, Quantification of ITG AV+ particles was performed on n ≥ 6 images acquired at 11,000 or 13,000X (approximately 100–700 particles per full image were analyzed). **a and c**, One-way ANOVA with Tukey’s multiple comparisons test. **e**, Kruskal– Wallis test with Dunn’s multiple comparisons test. Mean ± SD. *, **, ***, and **** represent *P* < 0.05, 0.01, 0.001, and 0.0001, respectively.

Beyond total ITG AV detection, we have performed immunogold labeling to detect surface ITG AV on the particles via TEM (**Fig. 3d**). Quantitative analyses showed a significantly higher percentage of particles with ITG AV surface detection in Mem-SVs, compared to Empty-SVs and Cyto-SVs. However, this trend was not significant in comparing Complete-SVs to Empty-SVs or Cyto-SVs. A possible explanation could be that the higher total protein loading during Complete-SV synthesis influences certain membrane protein incorporation efficiency. However, further investigation is warranted. In addition to ITG AV, we have employed single particle fluorescence imaging (SPFI), a recently developed fluorescence imaging approach for nanoparticles^33^, to confirm higher surface presence of fluorescently labeled CD63 on Mem-SVs and Complete-SVs than Empty-SVs and Cyto-SVs (**Supplementary Data 7b**).

Collectively, our application of multiple validation methodologies, from bulk detection to precise spatial localization, confirmed that a fraction of surface proteins successfully integrates or binds to SV membranes.

### Lipidomic Profiling Revealed Biological Lipids Enriched with Membrane Proteins

The extraction of proteins from native cells using detergents often accompanied by the co-extraction of lipids and residual detergent, resulting in the formation of lipid–protein–detergent complexes. These complexes are essential for the solubilization of membrane-associated proteins but can also introduce cellular (endogenous) lipids into downstream formulations. This lipid remodeling during SV synthesis represents a critical but underexplored aspect of biomimetic vesicle engineering. There has been limited investigation into the lipidome of cellular protein-loaded SVs, the relative abundance of endogenous lipids versus synthetic lipids, and the implications for future engineering of protein-lipid membrane mimetics.

To address this knowledge gap, we extracted lipids from MemRaw, CytoRaw, and SVs, and conducted liquid chromatography-tandem mass spectrometry (LC-MS/MS) in both positive (pos) and negative (neg) ionization modes (**Fig. 4a, Supplementary Data 8a-b**). Pos mode favors the detection of basic or neutral molecules such as phosphatidylcholine (PC), lysophosphatidylcholine (LPC), sphingomyelin (SM), ceramides (Cer), cholesterol esters (CE), diacylglycerols (DG), and triglycerides (TG), while neg mode enhances the detection of acidic lipids such as phosphatidylinositol (PI), phosphatidylglycerol (PG), phosphatidic acid (PA), phosphatidylserine (PS), and lysophospholipid^34^.

**Figure 4.**
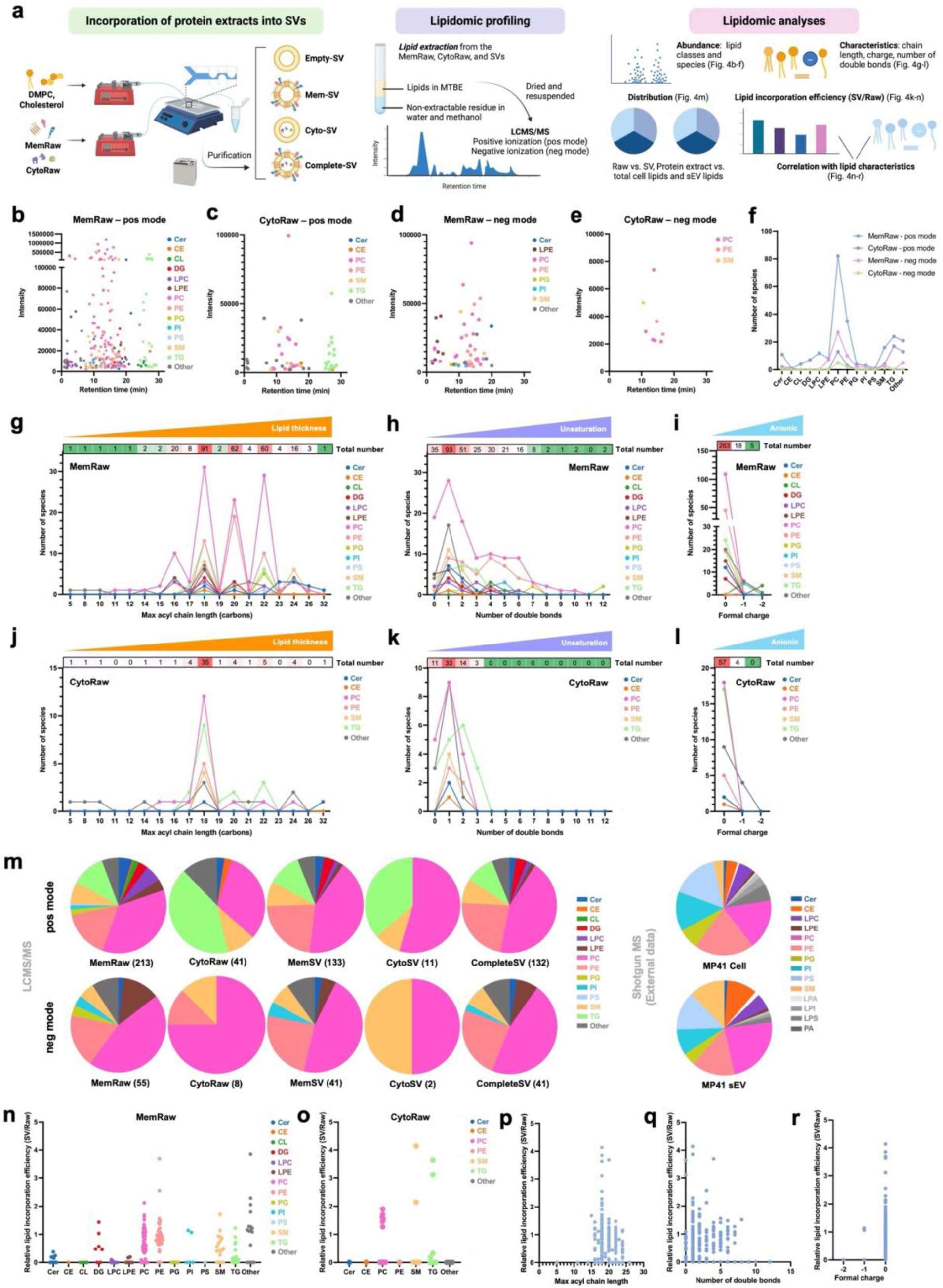
Lipidomic profiling of SVs and raw proteins. **a**, Schematic overview of sample derivation for lipidomic analysis. Summary of the lipidomic analyses performed. **b–e**, Retention time versus lipid intensity plots with color-annotated lipid classes for MemRaw in positive (pos) ionization mode (**b**) and negative (neg) mode (**d**), and for CytoRaw in pos (**c**) and neg (**e**) ionization modes. **f**, Number of lipid species detected in each lipid class. **g–l**, Lipid species grouped by maximum acyl chain length (**g, j**), number of double bonds (**h, k**), and formal charge (**i, l**) in MemRaw (**g–i**) and CytoRaw (**j–l**) samples, using data from both ionization modes. **m**, Lipid distribution profiles of raw proteins and protein-loaded SVs after removing confounding lipid annotations (mainly resulting from saturated DMPC peak in SVs) using Empty-SVs as a reference. External lipidomic profiles of MP41 cells and sEVs are shown^35^ for comparison. **n–o**, Relative lipid incorporation efficiency, presented as fold change in SVs versus corresponding raw proteins: MemSV/MemRaw (**n**) and CytoSV/CytoRaw (**o**) (combined both pos and neg modes). **p–r**, Correlation between lipid incorporation efficiency (SV/Raw) and lipid physicochemical properties, including maximum acyl chain length (**p**), number of double bonds (**q**), and formal charge (**r**), using data from MemSV and CytoSV across both ionization modes.

A wide lipid diversity was identified in the raw protein extracts. The majority of lipid species were identified in pos ionization mode, with 229 species detected in MemRaw and 53 in CytoRaw, complemented by 57 and 8 species detected in neg ionization mode, respectively (**Figs. 4b–e**). In MemRaw (pos mode), PC was the most abundant lipid class, comprising 35.81% of total lipids, followed by phosphatidylethanolamine (PE) (15.28%), TG (10.48%), SM (6.99%), lysophosphatidylcholine (LPC) (5.24%), ceramide (Cer) (4.80%), lysophosphatidylethanolamine (LPE) (3.49%), diacylglycerol (DG) (3.06%), cardiolipin (CL) (1.75%), PG (1.75%), PI (1.31%), cholesteryl ester (CE) (0.44%), and phosphatidylserine (PS) (0.44%). CytoRaw exhibited a markedly narrower lipid diversity and intensity compared to MemRaw (**Figs. 4b-f, Supplementary Data 8a–b**), indicating lower lipid complexity and abundance. The relative abundance of the detected endogenous lipids is generally lower than that of the synthetic DMPC and cholesterol, as visualized in chromatography (**Supplementary Data 8a-b**). The absolute amount of DMPC and cholesterol was determined by an external standard curve, and their molar ratio was determined to be approximately equal across SVs (**Supplementary Data 8c**).

To gain insights into the lipid environment in which endogenous proteins reside, we analyzed key lipid physicochemical properties, specifically bilayer thickness (represented by the number of carbons in the longest chain in a lipid), degree of unsaturation, and formal charge, in MemRaw and CytoRaw samples (**Figs. 4g–l**). Notably, while the maximum acyl chain length ranged from 5 to 32 carbons, 76% of all detected lipids in MemRaw exhibited chain lengths of either 18 (32.5%), 20 (22.14%), or 22 (21.43%) carbons. This enrichment was consistent across several major lipid classes, including PC, PE, and SM, with SM and Cer further enriched in longer chain lengths of 24 carbons. These results suggest that the membrane bilayer associated with membrane proteins predominantly features a thickness equivalent to 18–24 carbon chains (**Fig. 4g**). In CytoRaw, 57.38% of all detected lipids possessed an 18-carbon chain, whereas 6.56% and 8.20% of lipids had chains of 20 and 22 carbons, respectively (**Fig. 4j**), indicating a more homogeneous and thinner lipid environment associated with cytosolic proteins. In terms of saturation, monounsaturated lipids were the most prevalent for both MemRaw (32.52%) and CytoRaw (54.10%), followed by di-unsaturation (17.83% and 22.95%, respectively), saturation (12.23% and 18.03%, respectively), and higher unsaturation with individual species containing up to 12 double bonds (**Figs. 4h, 4k**). PC and PE were the major contributors to polyunsaturation (≥ 2 double bonds), underscoring their roles in maintaining membrane fluidity and integrity. Regarding formal charge, the majority of detected lipids, including phospholipids, SM, Cer, CE, and TG, were neutral. Anionic lipids such as CL, PI, PS, and PG, as well as fatty acids (categorized under “Other”) contributed to the negatively charged lipid population (**Figs. 4i, 4l**).

The lipid profile of MemRaw and CytoRaw remained largely preserved upon incorporation into SVs (**Fig. 4m**). To evaluate the biomimicry of SV lipid composition relative to native cellular and EV-derived membranes, we compared our data with the lipidomic profiles from total cellular lipids and sEVs derived from the same MP41 cell line^35^. In both MP41 cells and sEVs, PC, PE, and SM were the most enriched lipid classes, accompanied by notable levels of anionic lipids such as PI, PS, and PG, as well as CE and lysophospholipids (LPA, LPC, LPE, LPI, and LPS) (**Fig. 4m**). These lipids reflect the composition of typical mammalian plasma membranes and EVs, essential in organizing membrane microdomains (lipid rafts) (SM), membrane surface charge (PI, PS, and PG), signaling (SM, PI, PS, PG), and membrane curvature and fluidity (lysophospholipids)^36,37^.

Among the SV formulations, CompleteSVs were considered the most biomimetic because they incorporate membrane and cytosolic proteins together with lipids. Compared with the parental cells and natural EVs, CompleteSVs reproduced the major lipid classes, particularly PC, PE, SM, and PI, but also displayed distinct features, including the presence of CL and TG and the absence of lysophospholipid classes and PA. Together, these findings indicate partial rather than complete lipidomic mimicry. We note that this comparison is qualitative, given the different lipidomic workflows used for native cells/sEVs (shotgun lipidomics)^35^ and CompleteSVs (LCMS/MS). Furthermore, the protein and lipid extraction processes, together with vesicle reassembly during microfluidic synthesis, may not fully preserve the selective lipid composition of endogenous cells and EVs.

The finding that the lipidomic profiles of SVs were not fully identical to their corresponding raw protein extracts suggested a potential preferential loading. We therefore next investigated how lipid physicochemical properties are associated with incorporation efficiency— defined as the fold change of lipid abundance in SVs relative to corresponding lipids in the Raw proteins (SV/Raw). PC, PE, SM, DG, and TG exhibited higher incorporation efficiency overall compared to lysophospholipids and anionic lipids such as CL, PG, PI, and PS (**Figs. 4n–o**). This difference may be attributed to the intrinsic structural and biophysical characteristics of these lipid classes. Lysophospholipids, which contain only a single acyl chain^36^, may destabilize bilayer membranes and promote non-lamellar structures, possibly impairing their stable integration into vesicle membranes. In terms of bilayer thickness, lipids with maximum acyl chain lengths between 16 and 24 carbons were generally well-incorporated, with 18-carbon chains showing the highest efficiency (**Fig. 4p**). No clear correlation was observed between incorporation efficiency and the degree of unsaturation for lipids containing fewer than 8 double bonds (**Fig. 4q**). However, lipids with more than 8 double bonds appeared poorly incorporated, which could be associated with their increased structural complexity. Furthermore, lipids carrying a formal charge of −1 or −2 demonstrated markedly lower incorporation efficiency than neutral lipids (**Fig. 4r**), suggesting that electrostatic interaction could play a role in lipid incorporation during SV formation.

Taken together, these findings highlight that incorporating a biologically informed range of lipid classes, chain lengths, degrees of saturation, and charge states into SVs could be essential for recapitulating native membrane plasticity and structural stability, as well as compatibility with membrane-associated proteins.

### Membrane Protein Incorporation Enhanced Cell Uptake

Studying the respective roles of EV surface and internal proteins typically relies on genetic manipulation^5,8,38^, which could be limited to one or a few targets at a time and is constrained by the tolerance of the source cells. Our SV platform circumvents these limitations by enabling modular loading of membrane/cytosolic proteins - a unique advantage for dissecting compartment-specific contributions. A series of SVs was synthesized, including Empty-SVs, Mem-SVs, Cyto-SVs, Complete-SVs, and SVs loaded with varying amounts (1, 3, or 5 µg) of the integrin αVβ5 protein (designated as ITG AVB5-1/3/5UG-SVs) (**Fig. 5a**). Loading of recombinant ITG AVB5-loaded SVs was confirmed by SDS–PAGE (**Supplementary Data 7c**).

**Figure 5.**
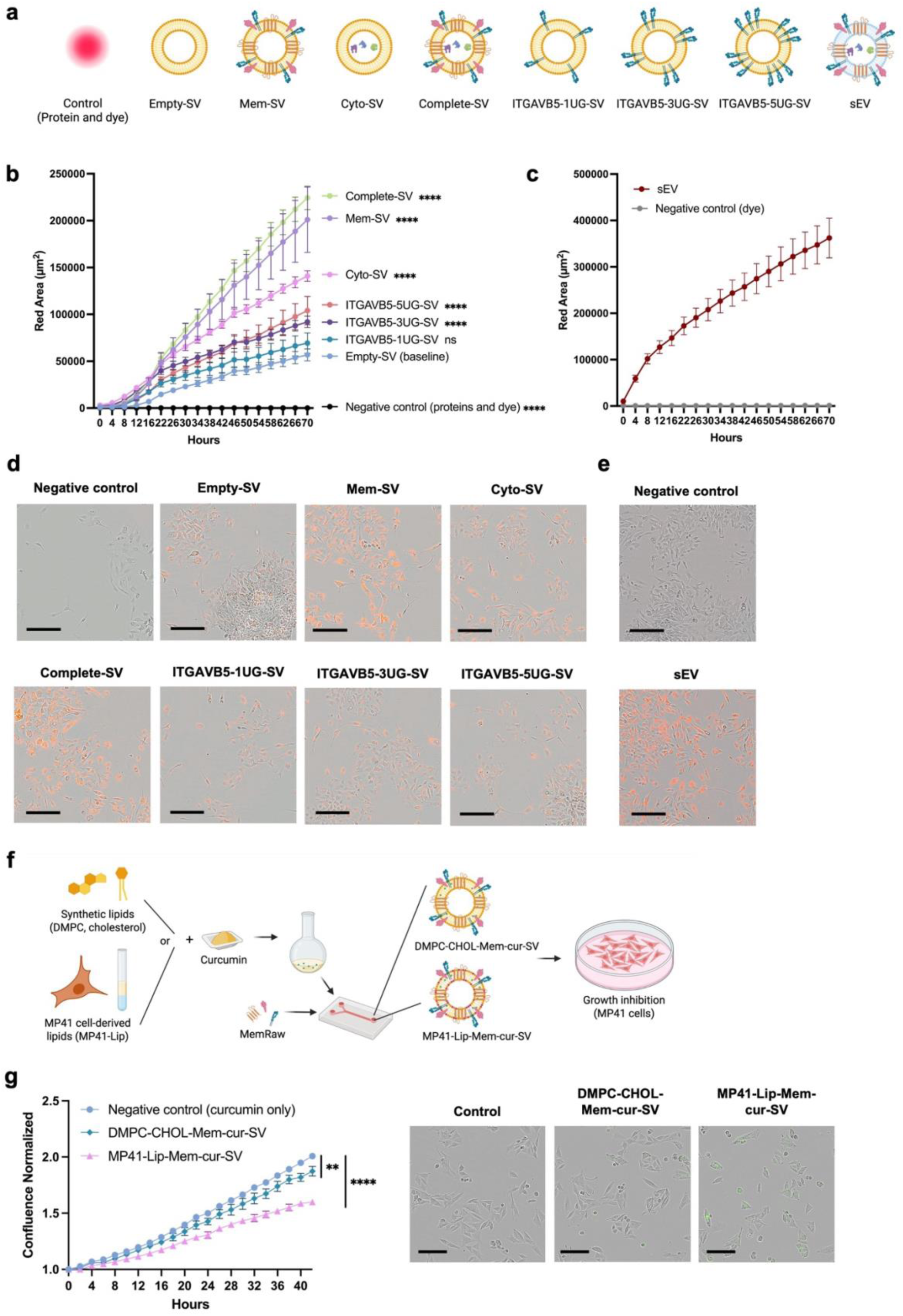
Selective protein loading and lipid optimization modulate cell uptake of SVs and the cytotoxic effect of curcumin delivery. **a**, Schematic for the different types of SVs synthesized, alongside sEVs, used for IHH (immortalized human hepatocytes) cell uptake comparison in **b-c**. **b-c**, IHH uptake of SVs treated at 1.3E10 particles/mL (**b**) and uptake of sEVs treated at E10 particles/mL (**c**), and their representative images at the endpoints were shown in (**d**) and (**e**), respectively. Scale bars = 200 µm. Red fluorescence: sp-DiIC18 dye-labeled particles. **b and d**, Negative controls for SV uptake represent blank syntheses without lipids but with an equivalent amount of buffer, proteins (membrane and cytosolic proteins), and stained with dye and purified in the same way. Negative controls for sEVs represent buffer-stained and purified in the same way. **f**, Schematics of synthesizing curcumin-loaded SVs using synthetic lipids (DMPC, cholesterol) or MP41 cell-derived lipids (MP41-Lip) for evaluating their effects on the growth inhibition of MP41 cells. **g**, Change in confluence of MP41 cells treated with curcumin-loaded SVs, represented as the ratio of confluence at each time point over the initial confluence. Representative images at 42 hours post-uptake were shown. Scale bar: 200 µm. Green fluorescence: curcumin. **b**, Two-way ANOVA with multiple comparisons versus the baseline group (Empty-SV), with Tukey’s correction. Significance at the final time point (70 h) is shown. **g**, Unpaired t-test. Mean ± SD. n = 4 experimental replicates. **, and **** represent *P* < 0.01 and 0.0001, respectively.

Prior *in vivo* studies investigating the role of cancer cell-derived EVs in premetastatic niche formation used intravenous EV doses equivalent to 5–10 µg of protein per injection^5,39–41^. In our system, 10 µg of MP41 sEV proteins corresponded to approximately 5.2E9 particles, which would correspond to an estimated initial circulating concentration of ∼1.3–2.6E9 particles/mL in a mouse, assuming dilution in ∼2 mL of blood volume. Based on this estimate, we tested *in vitro* uptake of EVs and SVs at E9 particles/mL to compare their uptake efficiency (**Supplementary Data 9a**). Among all vesicle types, sEVs exhibited the highest level of cellular uptake, followed by Mem-SVs and Complete-SVs. In contrast, the uptake of Empty-SVs, Cyto-SVs, and ITG AVB5-loaded SVs was minimal, suggesting that bulk membrane protein loading (MemRaw) has a more pronounced effect on enhancing uptake than cytosolic proteins or single-protein (ITG AVB5) loading.

The limited uptake efficiency of SV formulations with limited protein loading—Empty-SVs (no protein loading) and ITG AVB5-SVs (loaded only with recombinant ITG AVB5)— showed undetectable cellular uptake at E9 particles/mL, consistent with the generally lower cellular uptake efficiency of liposomes relative to EVs. Therefore, to further resolve differences in uptake among lower-performing SV types, a parallel experiment was conducted using a higher particle concentration at 1.3E10 particles/mL (**Figs. 5b, 5d**). Under these conditions, Complete-SVs and Mem-SVs consistently exhibited the highest levels of uptake, followed by Cyto-SVs, ITGAVB5-5UG-SV, and ITGAVB5-3UG-SV, which are significantly higher compared to Empty-SVs. Notably, the uptake of ITG AVB5-loaded SVs showed a dose-dependent trend based on the amount of protein loaded. SVs synthesized with 1, 3, and 5 µg of ITG AVB5 yielded red fluorescence areas of 6.9E4 ± 1.1E4 µm^2^, 9.2E4 ± 6.2E3 µm^2^, and 10.4E4 ± 1.5E4 µm^2^, respectively, at 70 hours post-treatment, indicating that ITG AVB5 plays a role in enhancing SV uptake by IHH cells. None of the SV types nor sEVs induced cytotoxicity in IHH cells at the tested concentrations (**Supplementary Data 9b**). As a comparison, sEV uptake was also tested at high concentration (E10 particles/mL) on IHH, which resulted in approaching 4E5 µm^2^ of red area, approximately double that of Complete-SVs and Mem-SVs (**Figs. 5c, 5e**). This finding suggests that protein loading, particularly membrane proteins, significantly enhanced SV uptake, but sEVs showed superior uptake efficiency than all formulations.

To support the application of SVs as EV-mimetic systems, we performed dose-dependent in vitro assays comparing Complete-SVs (EV-mimetic) and sEVs in stromal cells (**Supplementary Data 9c–g**). We found that at higher particle concentrations (E10 and 2E10 particles/mL), Complete-SVs significantly enhanced BJ fibroblast proliferation and did not significantly influence IHH cells, comparable to those observed with sEV treatment at lower particle concentrations (E8–E9 particles/mL). This observation is consistent with the substantially lower protein content per particle in Complete-SVs (approximately 50-fold lower than sEVs) (**Supplementary Data 9c**).

In addition to tuning protein composition, the lipid composition of SVs can be modulated to better mimic native membranes. To do this, we extracted total lipids from MP41 cells and used them to replace DMPC and cholesterol during SV synthesis. Furthermore, to demonstrate the therapeutic potential of the SV platform, we encapsulated a model hydrophobic drug, curcumin, into SVs, producing SVs with either synthetic lipids (Mem-cur-DMPC/CHOL-SVs) or MP41-derived lipids (Mem-cur-MP41Lip-SVs). Their effect on cancer cell growth was evaluated on the uveal melanoma cell line MP41 (**Fig. 5f**). Both SV formulations suppressed MP41 cell proliferation, with Mem-cur-MP41Lip-SVs exhibiting a more pronounced inhibitory effect (**Fig. 5g**).

To assess the broader adaptability of the SV platform, we produced vesicles using either cancer or non-cancer source materials (**Supplementary Data 10).** Compared with DMPC:CHOL-based vesicles, vesicles generated from natural lipid extracts (MP41, IHH, or HEK293T) were smaller and more negatively charged (**Supplementary Data 10a,b**). We further prepared four curcumin-loaded formulations using either synthetic lipids or HEK293T cell lipids, with or without HEK293T membrane proteins (HEK293T MemRaw) (**Supplementary Data 10c–f**). These formulations showed differential effects on MP41 cell proliferation. Specifically, HEK293T-Lip-Empty-cur-SVs resulted in complete inhibition of MP41 proliferation, followed by significant inhibition by HEK293T-Lip-Mem-cur-SVs and DMPC-CHOL-Mem-cur-SVs (**Supplementary Data 10d**). However, the proliferation-inhibitory effects of these SVs were not significant in IHH cells (**Supplementary Data 10e**), indicating selective toxicity toward MP41 cancer cells. The physicochemical properties and curcumin loading efficiency of these formulations suggest that the robust inhibition of MP41 cell proliferation may be associated with particle size-mediated cell uptake efficiency, the HEK293T lipid source, higher curcumin loading efficiency, or a combination of these factors. Taken together, these findings suggest that SVs may be applied for drug delivery applications with tunable protein and lipid profiles.

### Integrin Incorporation Modifies the Retention of SVs in Multiple Organs

Given that total cell membrane proteins, as well as specifically ITG AVB5, promote cellular uptake in IHH cells (**Fig. 5c**) and the association of ITG AVB5 in liver-tropism^5^, Mem-ITG AVB5-SVs were produced in addition to Empty-, Mem-, Cyto-, and Complete-SVs to investigate *in vivo* biodistribution (**Fig. 6a**). While the liver was the primary site of SV accumulation across all SV types, Mem-ITG AVB5-SVs showed significantly higher average radiant in the liver compared to all other SVs (**Fig. 6b**). Notably, Mem-ITG AVB5-SVs also showed significantly enhanced average radiant in the spleen and the lungs. Data has been plotted using different metrics, including total photons per organ and percentage of photons per organ over all measured organs. All metrics showed that Mem-ITG AVB5-SVs significantly enhanced liver accumulation (**Supplementary Data 11**).

**Figure 6.**
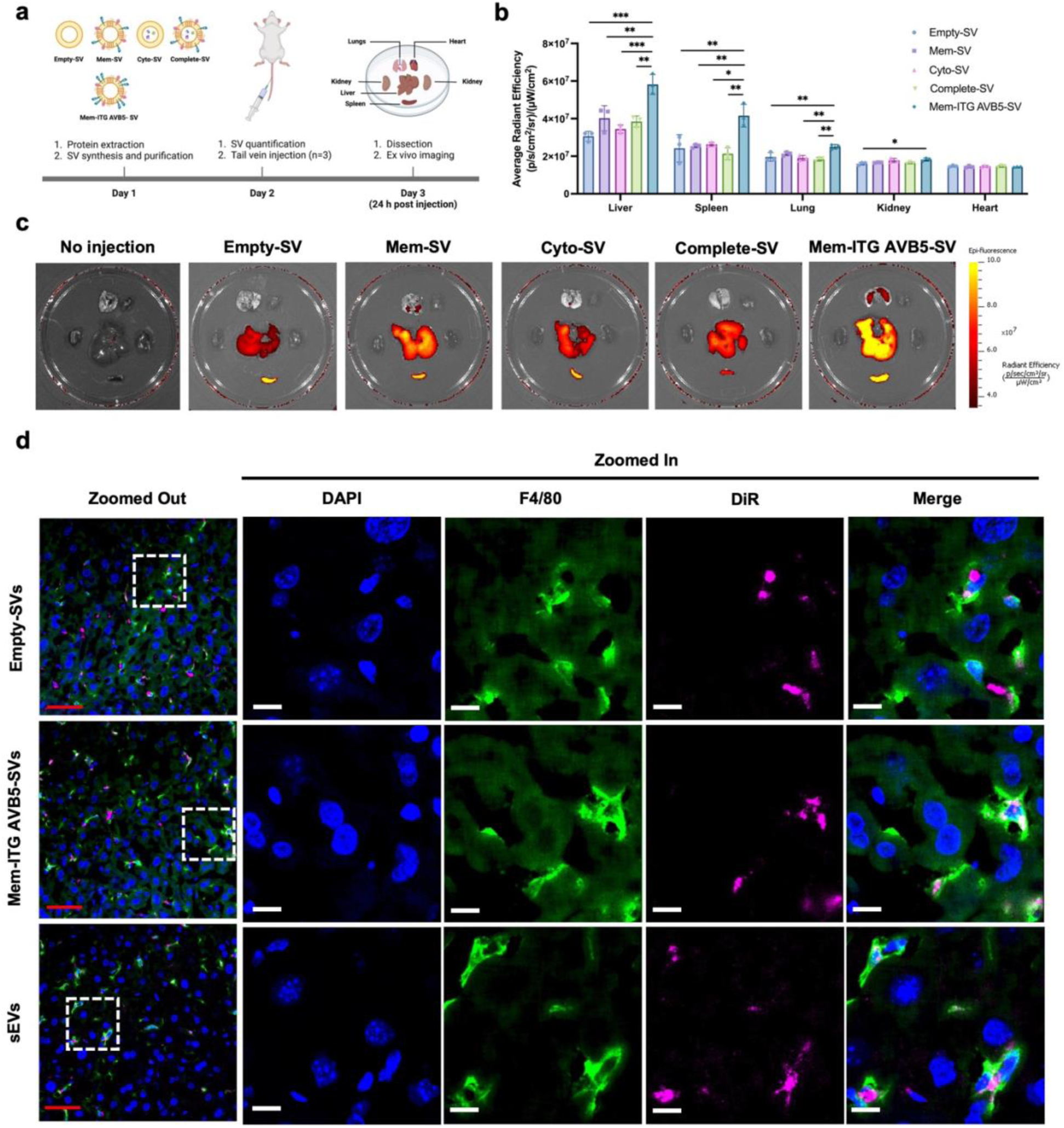
Biodistribution profile of SVs is influenced by protein incorporation. **a-c**, Comparison of biodistribution profiles of SVs by *ex vivo* imaging. SVs were labeled with DiIC18(5)-DS far red dye. All SVs were injected at 5E10 particles/mL. n = 3 mice per condition. **a**, Schematic of timeline for SV synthesis, tail vein injection, and *ex vivo* imaging. **b**, *Ex vivo* biodistribution of SVs, shown in average radiant efficiency. n = 3 mice. Fluorescence was quantified by Living Image® software. One-way ANOVA followed by Tukey’s multiple comparison test. Mean ± SD. *, **, and *** represent *P* < 0.05, 0.01, and 0.001, respectively. **c**, Representative *ex vivo* images of organs (lungs, heart, kidneys, liver, and spleen) collected at 24 h. **d,** Representative images of co-localization of Kupffer cells (F4/80+ cells in the liver) with SVs and sEVs. Red scale bars = 50 µm. White scale bars = 10 µm. Areas highlighted by a dotted square were zoomed in.

As the predominant tissue-resident macrophages, Kupffer cells play a central role in liver homeostasis, acting as scavengers with potent phagocytic activity^42^. Liver sinusoidal endothelial cells, a specialized endothelial subset, are crucial for hepatic immune regulation, serving as an interface for immune cell adhesion, including Kupffer cells^43^. Hoshino *et al.*^5^ reported that ITG AVB5-enriched exosomes co-localize with F4/80⁺ Kupffer cells in fibronectin-rich regions of the mouse liver. Based on this, we investigated whether SVs exhibit a similar co-localization pattern. We then sought to assess the localization of SVs with F4/80⁺ (**Fig. 6d**). We applied the phasor plot function for colocalization analysis (**Supplementary Data 12a-c**). A phasor plot transforms the fluorescence decay at each image pixel into a point on a two-dimensional map, where pixels with similar fluorescence lifetimes cluster together^44^. This allows tissue autofluorescence to be distinguished and filtered out from dye-associated fluorescence, which is especially useful in liver tissue where autofluorescence can spectrally overlap with fluorescent dyes and create apparent colocalization. SVs were detected co-localized with both F4/80⁺ Kupffer cells (**Fig. 6d**) and CD31⁺ endothelial cells (**Supplementary Data 12d**) in the mouse liver, indicating interactions with hepatic immune and vascular compartments.

Collectively, these findings highlight the potential of SVs as a model system to study vesicle-cell interactions and cell population dynamics within the tissue microenvironment at both the molecular and structural levels.

## Discussion and Conclusion

Following the discovery of EVs and the development of synthetic liposomes over the past half-century, a novel class of nanocarriers termed biomimetic SVs has emerged rapidly in recent years. These biomimetic SVs uniquely possess both the biocompatibility of EVs with the manufacturing advantages of synthetic liposomes^45^. Biomimetic SVs can be synthesized by a bottom-up approach, facilitating customizable composition, high scalability, and relatively high homogeneity. These advantages overcome the inherent heterogeneity and limited yield of natural EVs, thereby favoring clinical translation. Early research has focused on utilizing membrane proteins derived from leukocytes^18,19,46,47^, cancer cells^48,49^, and red blood cells^18,48^ to functionalize SVs. These SVs effectively replicate parental cell functionalities, such as tissue tropism and immune modulation, demonstrating therapeutic potential exceeding that of traditional EVs or liposomes alone.

Despite rapid advancements in biomimetic SV applications, research gaps remain in the technical development of quality control methods (*e.g.*, assessing sample purity, quantifying protein incorporation, determining protein localization), mechanistic insights into SV synthesis dynamics, the molecular composition, and the level of biomimicry. We introduce a robust SV platform to address these technical knowledge gaps, as well as to model cancer cell-derived EVs for dissecting EV-associated factors in intercellular communication. Critically, this work offers one of the earliest comprehensive proteomic and lipidomic comparisons between raw biomaterials and reconstituted SVs, and direct comparisons with EVs and synthetic liposomes in cell uptake. These analyses evaluated biomimicry levels in physicochemical properties, cellular uptake (*in vitro* and *in vivo*), elucidating the strengths and remaining challenges in transitioning from conventional EVs and liposomes to advanced biomimetic SVs.

Achieving efficient protein incorporation remains a major challenge and is underexplored in the development of protein delivery nanocarriers. A limited number of studies have investigated the encapsulation efficiency of individual or bulk proteins and explored strategies to enhance it by varying lipid formulations, production parameters, and synthesis methods^18,46,50^. Forbes *et al*.^50^ demonstrated that microfluidic-based liposome synthesis can achieve significantly higher protein loading (20–35%) compared to sonication or extrusion methods (<5%). Specifically, the incorporation efficiencies of insulin (5.7 kDa), ovalbumin (42.7 kDa), and bovine serum albumin (BSA, 65.5 kDa) were reported to be 36.8 ± 2.7%, 34.2 ± 4.9%, and 25.2 ± 2.8%, respectively^50^, suggesting that incorporation efficiency is protein type-dependent. Additionally, factors such as flow rate ratio (FRR) and total flow rate (TFR) have been shown to influence incorporation efficiency^18,46,50^. However, there is currently no consensus on a standardized set of parameters (*e.g.*, FRR, TFR, buffer composition, lipid formulation) for optimizing protein incorporation in the bottom-up synthesis of protein-functionalized liposomes^45^. Given the labor-intensive nature of assessing incorporation efficiency at one protein at a time, we present a systematic proteome-wide analysis using the input (raw) protein mixture and the encapsulated protein fraction, providing a comprehensive dataset for quantitative analyses. We identified protein loading as a non-uniform process influenced by characteristics, including molecular weight, charge, isoelectric point, post-translational modification (S-palmitoylation), structural characteristics (amphipathic helix, disorder degree), and stability are key determinants of protein incorporation efficiency.

Detecting protein incorporation is challenging, as no single method is sufficient to capture true protein conformation. To validate the presence of surface protein incorporation, we applied multiple complementary analytical approaches, including fluorescence-based surface detection and immunogold labeling, enabling both qualitative and quantitative visualization (**Fig. 3**). We confirmed that surface proteins, specifically ITG AV and CD63, are detectable on the surface of Mem-SVs and are present at higher levels compared to Cyto-SVs. However, the proportion of CD63-positive SVs in both Mem-SVs and Complete-SVs was approximately 50-fold lower than that observed in sEVs (**Supplementary Data 7b**). This relatively low surface protein detection in SVs may be attributed to differences in the cellular and EV proteome, as well as the substantially higher particle yield of SVs. The microfluidic synthesis method described in the present study achieved over 100-fold more particles per milliliter of production compared to MP41 sEVs isolated from 100 mL of conditioned medium in one minute of synthesis. Despite a range of EV-emission rates across cell types—0.18 EVs (erythrocytes) to 38 EVs (glioblastoma cells), 65 EVs (breast cancer cells), and 2,700 EVs (monocytes) per cell per hour^51–53^—biomimetic SVs enabled a much higher yield and efficiency at lower cost. However, the trade-off comes as requirements for a large protein amount to achieve protein densities comparable to those of sEVs. Thus, to enhance EV-mimetic protein density per particle, future work could optimize the synthesis parameters, such as increasing protein input or reducing the SV production yield. Future studies could quantify the proportion of membrane proteins, such as integrins, localized within the lumen versus on the surface of microfluidically generated SVs, to better define protein distribution, assess surface presentation efficiency, and inform subsequent functional studies. Furthermore, it should be noted that the lower protein density per SV may contribute to the observed reduction in subsequent cellular uptake efficiency (**Fig. 5**).

The lipid environment associated with cellular protein extract and the remodeling of the lipidome of SVs following cellular protein incorporation has been underexplored. Despite using fully synthetic lipids for all SV formulations, we report that the incorporation of protein extracts altered the overall lipidomic composition of the resulting liposomes. Therefore, it is likely that part of the changes observed in the physicochemical properties, cell uptake, and biodistribution could be attributed to the effects of lipids carried by proteins, despite their low abundance compared to the synthetic DMPC and cholesterol backbones (**Supplementary Data 8**).

The lipidomic profiles identified in the membrane protein extract revealed diverse lipid species and a distinctive profile compared to the native cell lipidome, offering novel insights for optimizing lipid formulations in biomimetic SV engineering. The broad distribution of acyl chain lengths and high degrees of unsaturation observed in MemRaw (**Fig. 4**) may reflect the biophysical requirements for embedding and stabilizing protein fragments or complexes within membrane-like environments^54^. In particular, long-chain and heterogeneous lipids could contribute to forming specialized microdomains or thicker bilayer regions that support integral membrane proteins and their conformational dynamics^36,55^. Furthermore, a deeper understanding of lipid sorting during EV biogenesis could further enable the rational design of vesicles.

A unique strength of SVs is their high customizability and flexibility for both basic biological research and the identification of candidate molecules for targeted therapeutic delivery. We loaded SVs with both bulk proteins (total cell membrane and/or cytosolic proteins) as well as individual commercial proteins (*e.g.*, recombinant ITG AVB5) to study cell uptake and biodistribution. This level of compositional control is difficult to achieve by EV engineering. We showed that membrane protein loading significantly enhanced cell uptake compared to cytosolic proteins and that loading ITG AVB5 moderately increased uptake, corroborating our initial hypothesis that among WCL, membrane proteins play the major role in mediating cell uptake. Interestingly, curcumin-loaded Mem-SVs synthesized with natural MP41 cell-derived lipids outperformed those generated with synthetic lipids (DMPC and cholesterol) in inhibition of cancer cell growth (**Fig. 5g**). This enhanced effect may relate to improved protein incorporation, drug loading efficiency, or synergistic interactions with natural lipids, though the mechanisms remain to be systematically investigated. Nonetheless, these findings highlight natural cell lipids as a promising parameter for optimizing the biomimetic and therapeutic potential of SVs.

Importantly, our study included negative controls consisting of protein-or dye-only, lipid-free syntheses to assess the removal of unbound proteins and dyes—an important aspect often overlooked in the literature. Although purification techniques such as dialysis and ultracentrifugation are commonly employed, they may not effectively eliminate all unbound proteins^56^ or free dyes^57^ due to the tendency of these molecules to aggregate. As a result, contaminants may be co-isolated with SVs, potentially leading to false-positive results in downstream analyses. The minimal protein bands observed in the protein-only control samples in our study support the successful removal of free proteins and lend greater confidence to our proteomic dataset and subsequent functional analyses related to cellular uptake and biodistribution.

A major challenge in tissue imaging, especially when translating *in vitro* findings to *in vivo* models, is tissue autofluorescence^58,59^. The use of the phasor plot to eliminate autofluorescence is critical in our study for minimizing false-positive signals during co-localization analysis, enabling more accurate visualization of SV interactions with hepatic immune and endothelial cells. We acknowledge that the biodistribution and macrophage colocalization results in the current study remain preliminary and do not establish functional biological changes. The contribution of this study is the fabrication of SVs and an assessment of EV-mimicry in proteomic and lipidomic profiles, cellular uptake, and biodistribution.

In summary, we present a rationally engineered SV platform that enables the systematic dissection of individual EV-associated factors, specifically membrane and cytosolic proteins, and their roles in cellular uptake and biodistribution. This bottom-up approach offers a versatile and controllable strategy to advance EV biology and serves as a blueprint for designing next-generation nanomedicines. By recapitulating key EV features in a simplified and tunable format, SVs function as “redesigned EVs” with tailored molecular composition and delivery potential.

Proteomic analyses in this study identified physicochemical features associated with efficient protein incorporation, while lipidomic profiling revealed the specific membrane environments that may support protein integration and functionality. Together, these omics data provide a molecular roadmap for optimizing biomimetic SVs. Moving forward, this platform can be adapted to model EV-mediated metastasis and other disease processes and enable the construction of functionally defined EV mimetics for precision delivery and therapeutic applications.

## Materials and Methods

### Cell Lines

MP41 (#CRL-3297) cell line was purchased from American Type Culture Collection (ATCC)^60^. Immortalized human hepatocytes (IHH) were gifted by Dr. Peter Metrakos (McGill University). 92.1 cells were kindly gifted from Dr. Martine Jager (Leiden University, Netherlands) ^61^. MP41 and 92.1 were cultured in RPMI-1640 (Corning^TM^ or Wisent Inc.) supplemented with 10% fetal bovine serum (FBS), 10 mM HEPES, 2 mM glutagro, 1 mM NaPyruvate, 1% penicillin and streptomycin (pen/strep) and 10 µg/ml insulin (Roche). IHH were cultured in DMEM with 4.5 g/L glucose, 10% FBS, and 1% pen/strep. IHH were cultured on poly-L-lysine (Millipore Sigma) coated surfaces. All cell lines were incubated at 37 °C with 5% CO2.

### Small Extracellular Vesicle (sEV) Isolation

Cells were cultured until they reached 70% confluency. Then cell culture medium was replaced by a medium supplemented with EV-depleted FBS (spun at 120,000 *g* for 18 h). 48 h after, the conditioned medium was collected, and cells were counted. The conditioned medium was subsequently centrifuged at 500 *g* for 20 min, 2000 *g* for 20 min, filtered (0.2 µm), and concentrated (Amicon Ultra centrifugal filters, 100 kDa cutoff). The concentrated medium was centrifuged at 120,000 *g* for 70 min in a TLA-110 rotor in Optima^TM^ MAX-XP ultracentrifuge. EV pellets were washed with filtered PBS, centrifuged at 120,000 *g* for 70 min, resuspended in filtered PBS, and stored at −80 °C. All centrifugation was done at 4°C.

### Protein Extraction and Processing

Cellular membrane and cytosolic proteins were extracted from MP41 separately using a Mem-PER^TM^ PLUS membrane protein extraction kit (Thermo Fisher Scientific). Briefly, cells were scraped in the culture medium, collected, and centrifuged to remove cell debris. Cells were counted to calculate the equivalent amount of reagent to be used based on the manufacturer’s protocol (0.75 mL of Permeabilization Buffer and 0.5 mL of Membrane Solubilization Buffer per 5 million cells). Cell pellets were washed twice using the Cell Wash Solution. Permeabilization Buffer and Solubilization Buffer were supplemented freshly with Halt^TM^ Protease and Phosphatase Inhibitor Cocktail (Thermo Fisher Scientific). Cell pellets were resuspended in the Permeabilization Buffer, and shaken on ice for 10 min, followed by centrifugation at 16,000 *g* for 15 min to collect the supernatant containing cytosolic proteins. Pellets were resuspended in the Membrane Solubilization Buffer, and shaken on ice for 30 min, followed by centrifugation at 16,000 *g* for 15 min to collect the supernatant containing membrane proteins. Proteins were used freshly within one day. Before use, the detergent was reduced from proteins by washing with 10 mL of MilliQ water three times (Amicon Ultra centrifugal filter, 10 kDa cutoff). Proteins were quantified using Pierce^TM^ Microplate BCA protein assay kit – reducing agent compatible. All centrifugation was done at 4°C.

### Synthetic Vesicle (SV) Synthesis and Purification

The lipid mixture was prepared freshly by mixing DMPC and cholesterol (each in chloroform form) at a 1:1 molar ratio. Chloroform was evaporated by a rotary evaporator and dried overnight in a vacuum desiccator. The dried lipid mixture was dissolved in 100% ethanol absolute, sonicated in a glass tube at 45 °C for 5 min, and filtered through a 0.2 µm PVDF filter. The lipid mixture in ethanol was stored at −20 °C for up to one month. Before use, the lipid mixture was adjusted to 9 mM with filtered ethanol and sonicated at 45 °C for 5 min. Proteins in MilliQ water were prepared based on the protein-to-lipid weight ratio (1:20 for Mem-SVs and Cyto-SVs, 1:10 (combination of 1:20 of membrane and cytosolic proteins, respectively) for Complete-SVs) and warmed up at 42 °C for 1 min. SVs were synthesized using a 3D-printed microfluidic chip^62^ on a heat plate (42 °C). Flow rate ratio (FRR) (aqueous phase to lipid phase) = 2 and total flow rate (TFR) = 60 mL/h according to Zinger *et al.*^19^. The microfluidic chip was rinsed with Ethanol and MilliQ water at FRR = 1 and TFR = 80 mL/h for 6 min before and after all syntheses, and 2 min in between syntheses. After syntheses, SVs were purified by ultracentrifugation at 120,000 *g* for 70 min, washed with an equal volume of MilliQ water, and spun at 120,000 *g* for 70 min. SVs were stored at 4 °C and used for experiments within three days.

### Characterization of Physical Properties

sEVs and SVs were characterized by ZetaView for size, concentration, and zeta potential. Their morphology was observed by TEM. For TEM, 10 µl of EVs or SVs were deposited onto negative glow-discharged carbon film 200 Mesh cuTEM grids (agar Scientific) for 30 min. The grids were washed with 3 drops of glycine (if fixatives are added), followed by 4 drops of water (2 min per wash). The grids were stained with 2% uranyl acetate for 1 min. The grids were dried overnight and then observed with a FEI Tecnai G2 Spirit Twin 120 kV Cryo-TEM.

### SDS-PAGE Stain-Free Gel Detection and Western Blot

SVs or EVs were mixed with Laemmli buffer and 2-Mercaptoethanol and heated at 95 °C for 10 min, followed by loading onto 4% −12% Mini-PROTEAN® TGX Stain-Free Protein Gels (#4568094, Bio-Rad). Protein bands in the gel were visualized using the ChemiDoc^TM^ XRS+ System (Bio-Rad). Proteins were then transferred onto polyvinylidene fluoride membranes (Bio-Rad). Membranes were blocked for 1 h in 5% non-fat dry milk in Tris-buffered saline with 0.1% Tween-20 (TBST). The target proteins were probed with primary antibody for anti-Integrin alpha V (1:1000, #ab179475, abcam), CD81 (1:1000, #ab109201, abcam), HSP90 (1:1000, #ab13492, abcam), TSG101 (1:1000, #ab125011, abcam), actin (1:1000, #A2229, Sigma-Aldrich), or syntenin-1 (1:1000, #ab19903, abcam) overnight. Membranes were washed in TBST, treated with secondary anti-rabbit antibody (1:1000, #7074S, Cell Signaling) or anti-mouse antibody (1:1000, #1706516, Biorad) in 5% milk for 1 h. Membranes were washed, developed using ECL prime Western blot detection (GE HealthCare), and visualized using the ChemiDoc^TM^ XRS+ System.

### Fluorescence Labeling and Purification for Vesicles

sEVs and SVs were labeled by sp-DiIC18 (#D7777) for *in vitro* cell uptake experiments, and DiIC_18_(5)-DS (#D12730) or DiR (#D12731) for biodistribution experiments (all from Thermo Fisher Scientific). Labeling was done based on a ‘salt-change’ method described by Cha *et al.*^57^. Briefly, EVs were washed twice with MilliQ water to minimize PBS concentration (Amicon Ultra centrifugal filter, 100 kDa cutoff), as PBS induces aggregations of the dyes^57^. EVs and SVs were stained with 10 µM of the dye at 37 °C for 30 min. PBS was then added to increase the volume by 10 times to induce aggregation of excessive dyes. Samples were incubated in PBS at room temperature for 10 min, followed by filtering (0.2 µm) and concentrating (Amicon Ultra centrifugal filter, 100 kDa cutoff). A negative control (mock liposome synthesis without adding lipids but protein only) was stained in the same way to validate the effectiveness of removing free dyes. Stained EV/SVs were used freshly within one day.

### Live-Cell Tracking of Cell Uptake of Vesicles

IHHs were seeded in a 96-well plate at 2500 cells/well 24-48 h before EV or SV treatment. Following quantification of stained sEVs and SVs by ZetaView, the concentration of sEVs and SVs was adjusted and controlled for EV and SV treatment. Briefly, cell culture medium was removed and replaced with a fresh complete medium supplemented with an equal number of particles in an equal composition of buffer (MilliQ and PBS) volume. The buffer volume was restricted to 10% of the total medium volume to account for the effect of the buffer on cells. Proteins and dye-only control were included. Cells were imaged by the IncuCyte S3 Live Cell Analysis Instrument. Four 10X images were taken in each well at an interval of 2 h. Cell confluence and fluorescence intensity were analyzed using IncuCyte S3 2021A Software.

### Single-Particle Protein Characterization

Single particle fluorescence was analyzed by CytoFLEX (Beckman Coulter) to characterize particles labeled with ITG AVB5 by Alexa Fluor 488 anti-human/rat CD51 (Integrin avb5) antibody (#920009, BioLegend). Size exclusion (qEV Gen 2, Izon) was performed according to the manufacturer’s protocols to remove unbound antibodies. For immunogold labeling, EVs and SVs were deposited on grids as described above and washed. Samples were blocked in 1% BSA (in PBS) for 5 min, then incubated with primary antibody (Integrin alpha V (1:1000, #ab179475, abcam)) at a 1:20 dilution at room temperature for 1 h. Samples were then washed with five drops of filtered DPBS for 5 minutes each, blocked in 1% BSA for 5 min, then incubated with secondary antibody (Goat anti-rabbit IgG H&L (10 nm Gold) (#ab27234, abcam)) at a 1:20 dilution at room temperature for 1 h. Samples were washed with DPBS and then stained with 2% uranyl acetate, as aforementioned. The grids were dried overnight and then observed with a FEI Tecnai G2 Spirit Twin 120 kV Cryo-TEM. Probing for CD63 was performed by size photometry and fluorescence imaging, following the methods reported by Wallucks *et al.*^33^.

### Mass Spectrometry for Proteomic Analyses

SVs were synthesized and purified as described (n = 3 syntheses). The raw proteins (post-detergent reduction, in MilliQ water) used for each synthesis were used as a reference (n = 3 protein extracts). To SV pellets post ultracentrifugation and to raw proteins at equivalent amounts, 400 µL RIPA buffer (Thermo Fisher Scientific) supplemented with Halt^TM^ Protease and Phosphatase Inhibitor Cocktail was added. Lysates were shaken on ice for 30 min, followed by the addition of 100 µL of 77% trichloroacetic acid (#T4885, Sigma-Aldrich) and shaken on ice for 1 h for protein precipitation. Samples were spun at 20,000 *g* for 20 min at 4 °C. The supernatant was discarded. Pellets were resuspended in 500 µL 90% cold acetone and incubated overnight. Samples were spun at 20,000 *g* for 20 min at 4 °C. The supernatant was removed and the pellet was dried at room temperature for 1 min. The pellet was then resuspended in 70 µL of Ammonium Bicarbonate for protein quantification and mass spectrometry. For each sample, proteins were loaded onto a single stacking gel band to remove lipids, detergents, and salts. The gel band was reduced with DTT, alkylated with iodoacetic acid, and digested with trypsin. Extracted peptides were re-solubilized in 0.1% aqueous formic acid and loaded onto a Thermo Acclaim Pepmap (Thermo, 75uM ID X 2cm C18 3uM beads) precolumn and then onto an Acclaim Pepmap Easyspray (Thermo, 75uM X 15cm with 2uM C18 beads) analytical column separation using a Dionex Ultimate 3000 uHPLC at 250 nl/min with a gradient of 2-35% organic (0.1% formic acid in acetonitrile) over 3 hours. Peptides were analyzed using a Thermo Orbitrap Fusion mass spectrometer operating at 120,000 resolution (FWHM in MS1) with HCD sequencing (15,000 resolution) at top speed for all peptides with a charge of 2+ or greater. The raw data were converted into *.mgf format (Mascot generic format) for searching using the Mascot 2.6.2 search engine (Matrix Science) against human protein sequences (Uniprot 2024). The database search results were loaded onto Scaffold Q+ Scaffold_5.0.1 (Proteome Sciences) for statistical treatment. *P* value and fold change were analyzed by Scaffold 5. Enrichment analysis for cellular components was analyzed by gene ontology terms (DAVID Database^63^). Protein sequence properties were analyzed by Multiple Protein Profiler^64^ for physicochemical properties, Protein-Sol^65^ for solubility, GPS-Palm^66^ for S-palmitoylation prediction, HeliQuest^67^ for amphipathic helix prediction, and IUPred2A^68^ for disorder score. For HeliQuest analysis, the helix with the highest hydrophobic moment per protein was selected, provided it met the following criteria: a hydrophobic moment > 0.4, a net charge > –1, and a hydrophobic face consisting of 5–9 amino acids.

### Lipid Extraction

Lipids from MemRaw, CytoRaw, Empty-SVs, Mem-SVs, Cyto-SVs, and Complete-SVs were extracted using the MTBE extraction method^69^. All SVs were synthesized using a 3 mM total lipid concentration of a DMPC and cholesterol mixture and were purified by dual ultracentrifugation, as outlined in the SV synthesis methods. For LC-MS/MS experiments, 100 µL of MemRaw (533 µg/mL), 200 µL of CytoRaw (465.5 µg/mL), and 300 µL of each SV sample were used. Prior to extraction, 5 µL of standard lipid mix (SPLASH™ LIPIDOMIX™ Mass Spec Standard, Avanti) was added to each sample. For every 100 µL of sample, 230 µL of methanol was added and vortexed, followed by the addition of 770 µL of MTBE. Samples were then shaken at room temperature for 1 h. Subsequently, 280 µL of MilliQ water was added, and the samples were vortexed and incubated at room temperature for 10 min. Phase separation was achieved by centrifugation at 1000 *g* for 10 minutes. The upper MTBE phase containing lipids was carefully collected and dried using a vacuum centrifuge. Dried lipids were then resuspended in 50 µL of isopropanol for downstream analysis.

### Lipid Identification with Liquid Chromatography-Mass Spectrometry (LC-MS/MS)

LC-MS/MS analyses were collected at the Proteomics and Molecular Analysis Platform, Research Institute of McGill University Health Centre. All reagents and solvents were LC-MS grade. OmniSolv® acetonitrile (ACN) and methanol (MeOH) were sourced from Millipore Sigma (St. Louis, MO, USA), Optima™ isopropanol and formic acid from Fisher (Fisher Chemical™), and Emsure® Ammonium acetate from Millipore (Millipore MOA, USA). Ultra-pure deionized water was obtained from the Millipore Milli-Q water system (Millipore MOA, USA). LC-MS analyses were performed with an ESI-QTOF-MS system (Impact II, Bruker, Billerica, MA, USA) coupled with a liquid chromatography system (1290 Infinity, Agilent) and equipped with a reversed-phase Agilent InfinityLab Poroshell 120 EC-C18 column (2.1 x 75 mm, 2.7 μm). Mobile phases were acetonitrile: water (40:60) at 10 mM Ammonium Formate with 0.1% Formic acid (A) isopropanol: acetonitrile (9:1) at 10 mM Ammonium Formate with 0.1% Formic acid (B). The following gradient elution method was used: 40 to 50% B from 0 to 2 min, 50−60% B from 2 to 3 min, a 60% B plateau for 3 to 8 min, 60−78% B from 8 to 18 min, followed by 78 to 99% from 18 to 31 min and a 99% B plateau for 5 min and a return to initial conditions with equilibration for 4 min. The column temperature was set at 50 °C, and the flow rate was 0.4 mL/min. The samples were kept in an autosampler at 10 °C. Each sample was injected at a volume of 10 μL. The operating parameters of the mass spectrometer were as follows: positive/negative spray voltage at 4500/3500 V, dry temperature at 230 °C, dry gas flow at 8 L/min, and nebulizer at 2 bar. The data were collected with a mass range from 50 to 1500 m/z with an acquisition rate of 8 Hz and stepping. The Auto MS/MS mode was used to obtain fragment ions. The instrument was calibrated using a 10 mM sodium formate solution before the runs, and, at the beginning of each run, this solution was injected as an internal calibrant for MS spectrum calibration. The LC-MS acquired data were processed using DataAnalysis Version 5.2 software (Bruker) and Bruker Compass Metaboscape 2023b software (Version 11.0.4, Bruker Daltonics GmbH & Co. KG).

### Biodistribution

Animal experiments were carried out under the approved animal protocol (MUHC-AUP#10108). Fluorescence-labeled SVs and sEVs were quantified by ZetaView. The concentration of particles was controlled with a final volume of 100 µL. SVs and sEVs were injected into NSG mice (< one year old, all males or all females in each experiment) intravenously (n=3 mice per type of SVs). After 24 hours, mice were sacrificed. Lungs, heart, liver, kidneys, and spleen were harvested and imaged by the Xenogen IVIS system.

Tissue processing and immunofluorescence. Harvested organs were fixed overnight with 4% PFA, followed by overnight submerging in 15% and another overnight submerging in 30% sucrose in PBS. Fixed tissues were then embedded with the O.C.T freezing reagent (Tissue-Tek®) and stored in −80 °C. 7 μm O.C.T tissue cryosections were made and stored in −20 °C. For immunofluorescence, cryosections were thawed and warmed up at room temperature for 10 min. Tissue sections were blocked using 1% BSA in PBS for 1 h. Primary antibodies were diluted in the blocking buffer: F4/80 (1:50, eBioscience; BM8), CD31 (1:100, Santa Cruz; MEC 13.3). Secondary antibodies conjugated to Alexa Fluor 488 or 549 were used (A-11001 and A-11007, Life Technologies). Fluorescent images were obtained by confocal microscope (Leica STELLARIS 8-STED-FLIM) and analyzed using LAS X Life Science Microscope Software.

### Statistical Analyses

EV isolation, SV syntheses, physicochemical characterizations, and cell uptake experiments were repeated at least three times independently. Comparison across different types of SVs and EVs was done by One-way ANOVA followed by Tukey’s multiple comparison test or unpaired t-test (specified in each Figure caption). *P* < 0.05 was considered significant.

## Acknowledgment

We sincerely thank the Centre for Applied Nanomedicine (CAN) at the RI-MUHC, led by Dr. Janusz Rak, for their support. We are grateful to Dr. Nadim Tawil and Laura Montermini for their assistance with training and experiments. Special thanks to Dr. Brian Meehan for IVIS instrument training for animal imaging and to Dr. Hong Choi for supporting the lipidomic analysis. We are thankful for the McGill Facility for Electron Microscopy Research, led by Dr. Kelly Sears, and to Jeannie Mui for training in TEM and immunogold labeling. We also appreciate the support of the Proteomics and Molecular Analysis Platform at RI-MUHC, led by Lorne Taylor and his team, Amy Wang and Jennifer Nedow, for conducting mass spectrometry experiments for proteomic analyses. Additionally, we acknowledge Dr. Joel Ryan at the McGill University Advanced BioImaging Facility (ABIF), RRID:SCR_017697, for his assistance with tissue image acquisition and analysis. We are thankful for the training on the use of CytoFLEX by Hélène Pagé-Veillett from the Immunophenotyping platform at the RI-MUHC. We are thankful for the training on the use of SPFI by Dr. Andreas Wallucks and Molly Shen from Dr. David Juncker’s laboratory. We also acknowledge Dr. Ruben Lopez and Chaymaa Zouggari for their training in the use of syringe pumps and the 3D printer (CADworks3D). Lastly, we appreciate our current and past lab members, particularly Tad Wu, Ishman Kaur, and Armen Erzingatzian, for their insightful discussions.

## Supplementary Data

The raw data that support the results are summarized in the supplementary data files or are available from the corresponding author upon request.

**Supplementary Data 1.**
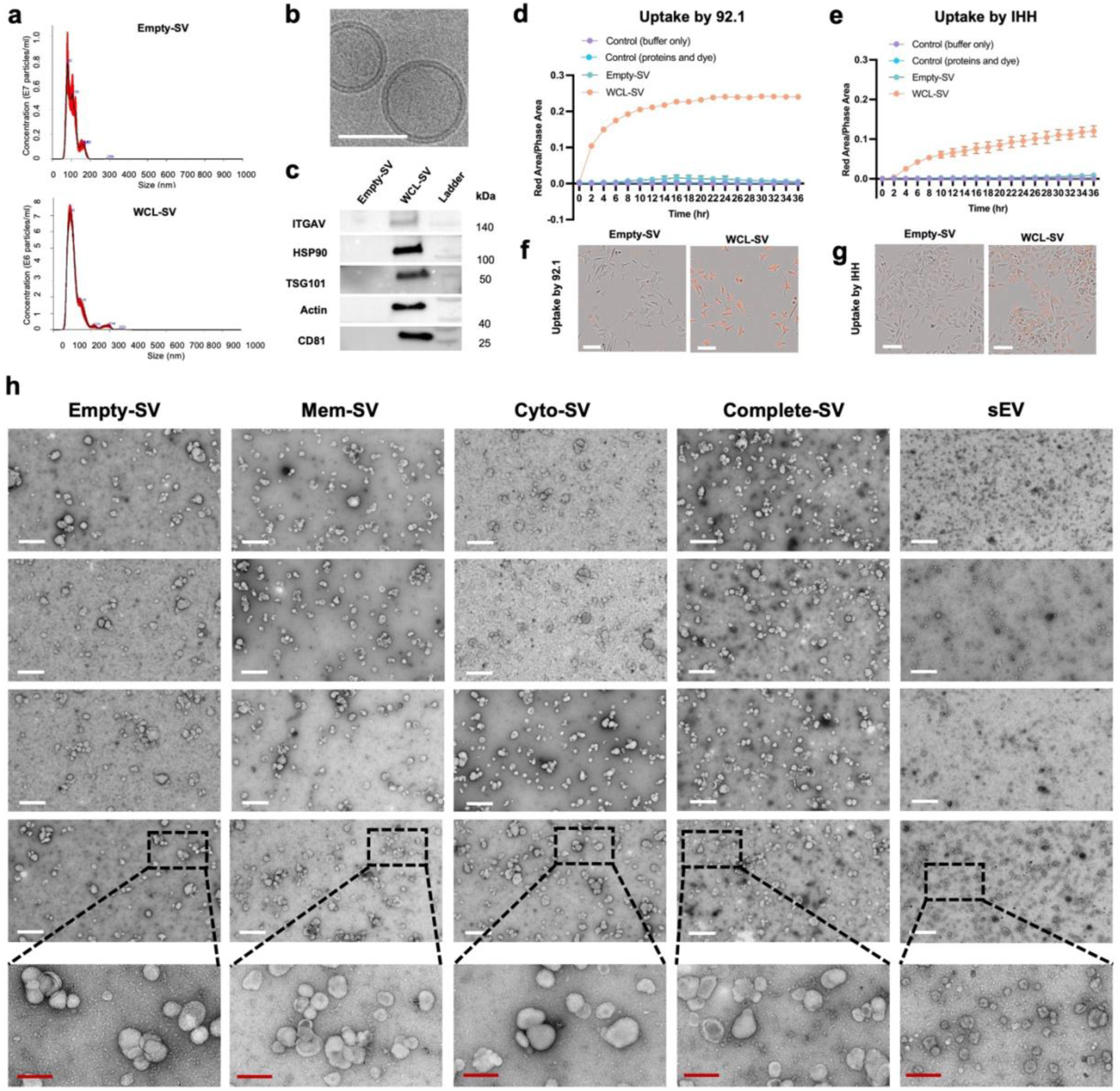
Supplementary characterization of SVs. **a**, Size distribution of Empty-SVs (no protein loading) (top) and WCL-SVs (loaded with WCL) (bottom) measured by NanoSight. **b,** Lipid bilayered membrane of Empty-SVs imaged by Cryo-TEM. Scale bars = 50 nm. **c,** Selected proteins of interest were detected in WCL-SV. Uptake of Empty-SVs and WCL-SVs (2.5E9 particles/mL) by uveal melanoma cells (92.1) (**d**) and immortalized hepatocytes (IHHs) (**e**) over 36 hours. Mean ± SD. Representative images at 24 hours for 92.1 cells (**f**) and IHHs (**g**). (**f-g**) Scale bars = 100 µm. **h**, Morphology of SVs loaded with MP41 cell membrane (Mem-SV), cytosolic (Cyto-SV), or both membrane and cytosolic proteins (Complete-SV), imaged by TEM. White scale bars = 600 nm. Red scale bars = 200 nm.

**Supplementary Data 2.**
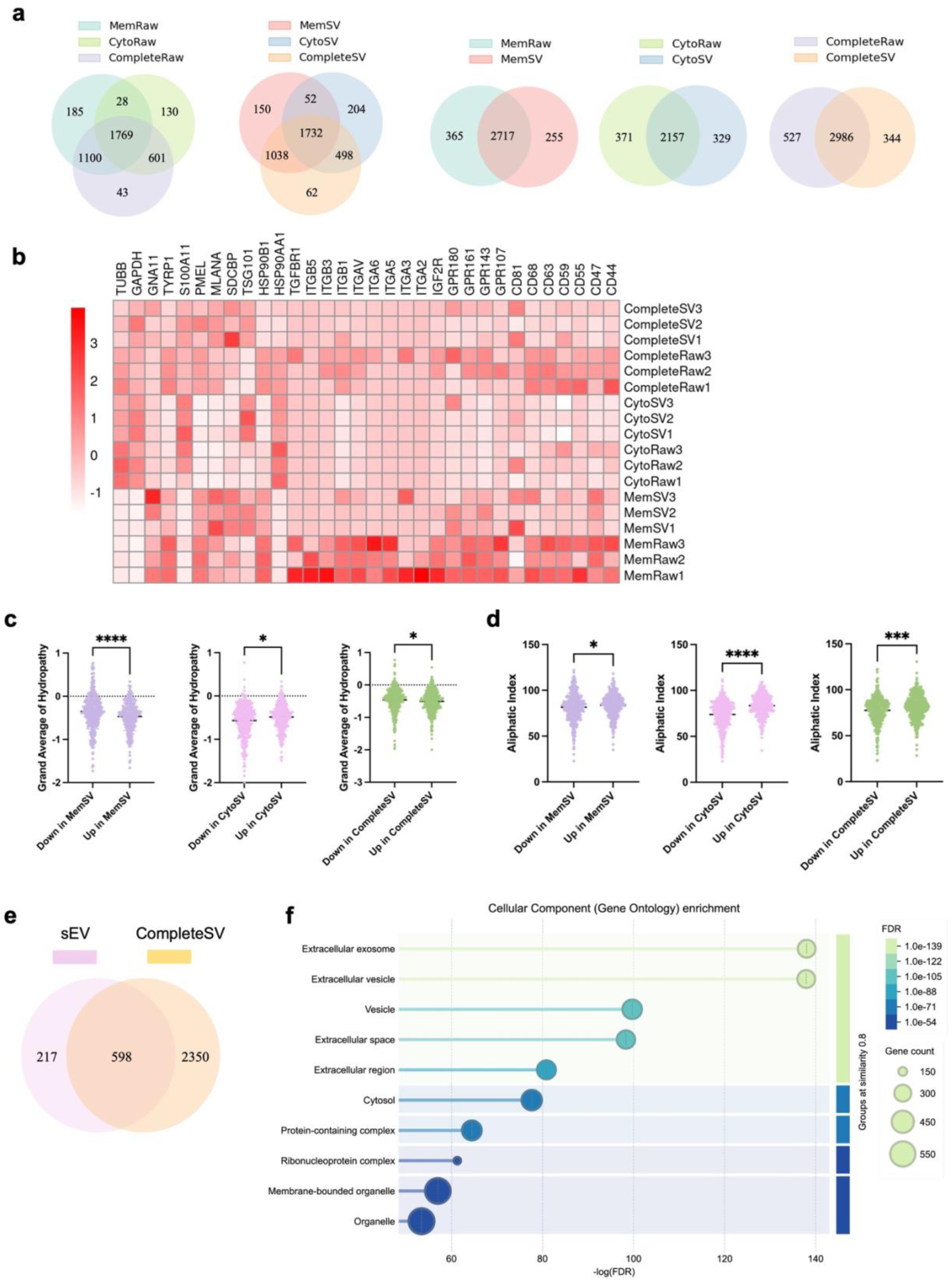
Extended proteomic data. **a**, Venn diagrams illustrating the overlapping and unique proteins (identified in at least one replicate) in each type of SVs and raw proteins. **b**, Heatmap of the abundance of proteins of interest. Comparisons of proteins in Down and Up in MemSV/CytoSV/CompleteSV for their protein grand average of hydropathy (**c**) and aliphatic index (**d**). **e**, Venn diagram showing the overlap between proteins identified in MP41 sEVs and CompleteSVs. **f**, Cellular component Gene Ontology enrichment analysis of the overlapping protein set between MP41 sEVs and CompleteSVs. **c-d**, Comparison of the protein physicochemical properties of significantly preferentially (Up) and poorly (Down) incorporated. Mann–Whitney U test.

**Supplementary Data 3.**
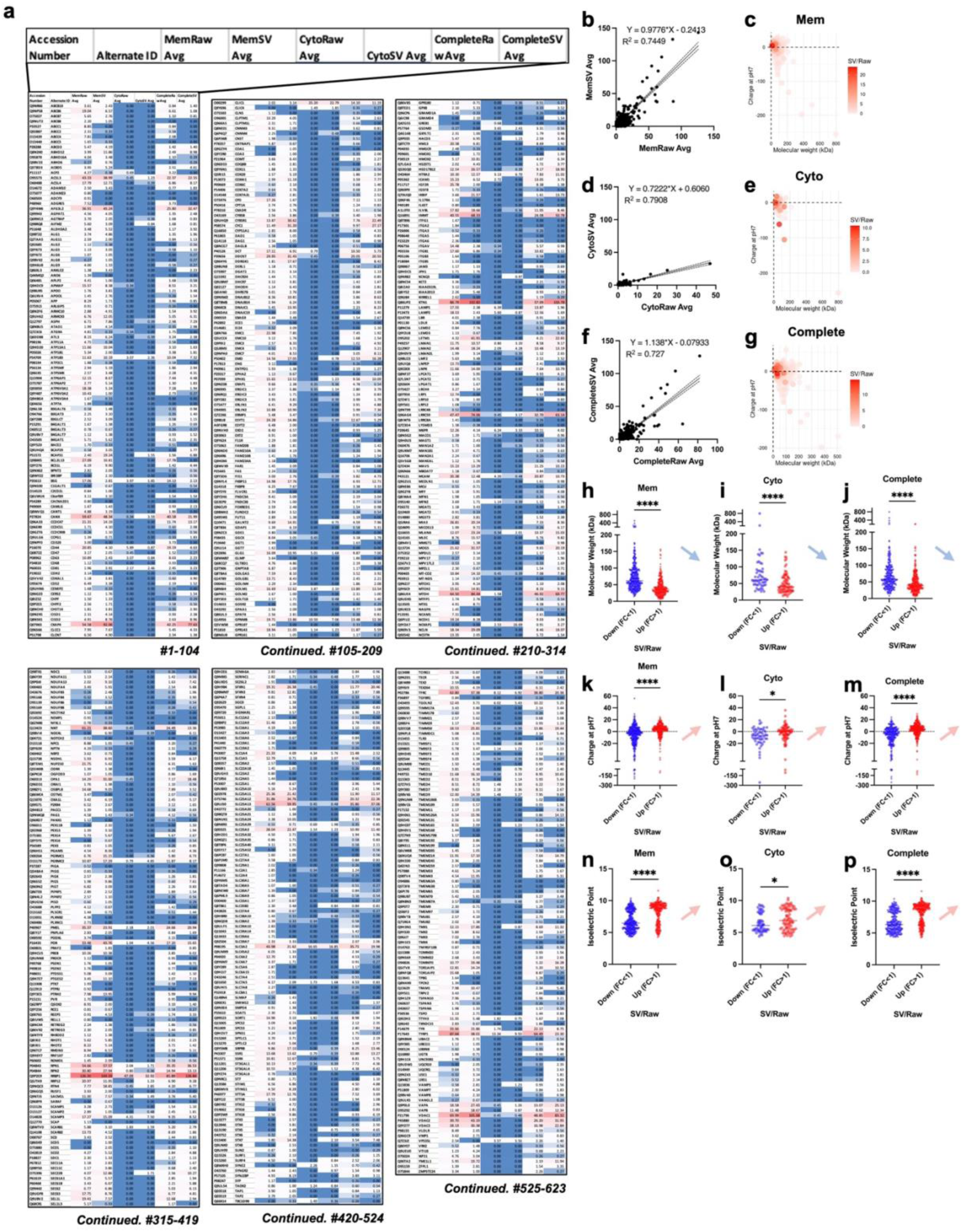
Transmembrane protein identification and analyses of incorporation efficiency. **a**, List of proteins with transmembrane localization, annotated using UniProt database, and their abundance (average spectra counts) in raw proteins and SVs. Correlations between the average spectra counts in MemRaw *versus* MemSV (**b**), CytoRaw *versus* CytoSV (**d**), and CompleteRaw *versus* CompleteSV (**f**). Correlations between charge, molecular weight, and protein incorporation efficiency (SV/Raw) (**c**, **e, g**), respectively. Molecular weight (**h-j**), charge at pH7 (**k-m**), and isoelectric point (**n-p**) of the poorly incorporated (Down, SV/raw (FC) < 1) and highly incorporated (Up, SV/raw (FC) > 1) proteins.

**Supplementary Data 4.**
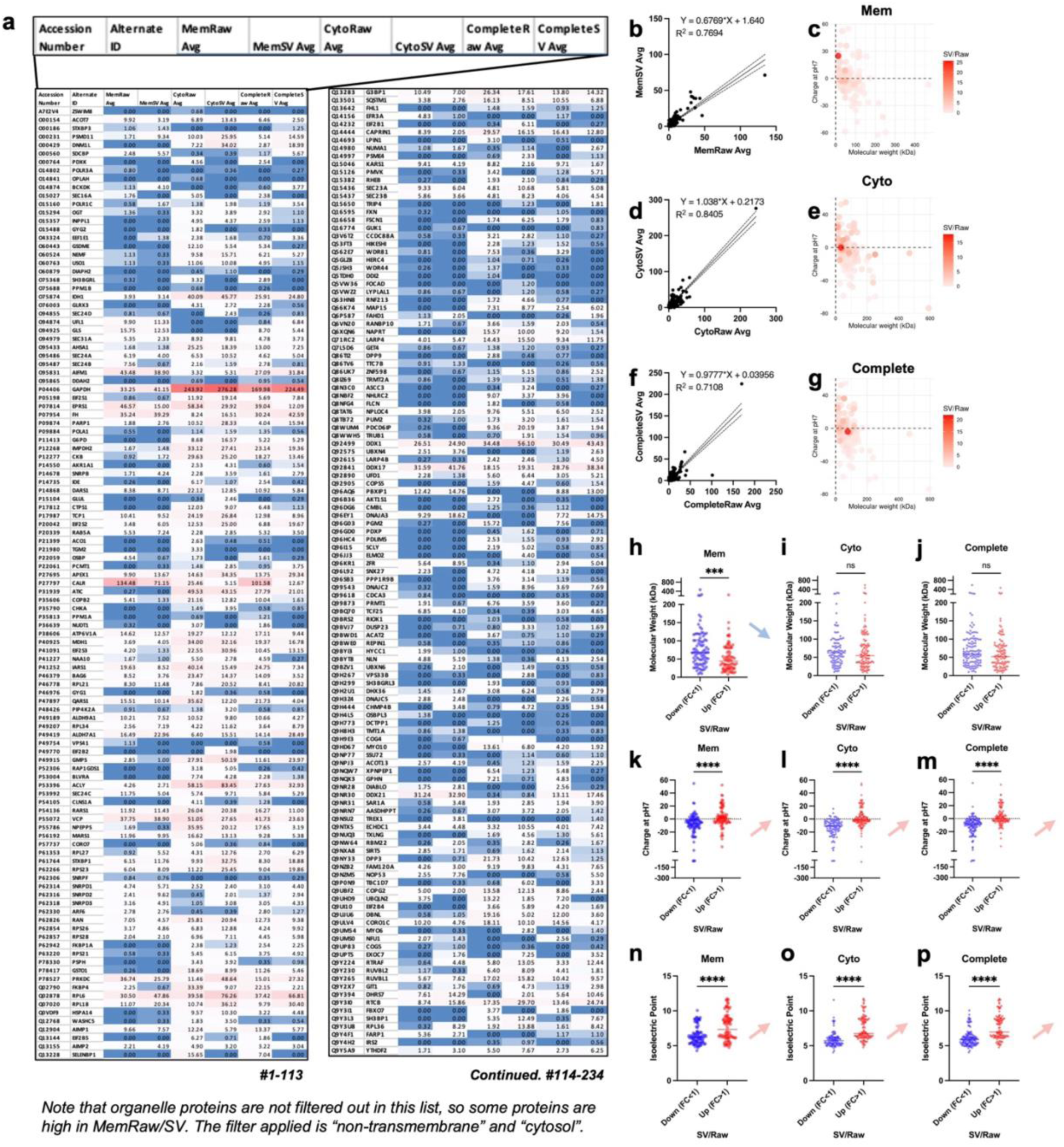
Cytosol and nontransmembrane protein identification and analyses of incorporation efficiency. **a**, List of proteins with cytosol but nontransmembrane localization, annotated using UniProt database, and their abundance (average spectra counts) in raw proteins and SVs. Correlations between the average spectra counts in MemRaw *versus* MemSV (**b**), CytoRaw *versus* CytoSV (**d**), and CompleteRaw *versus* CompleteSV (**f**). Correlations between charge, molecular weight, and protein incorporation efficiency (SV/Raw) (**c**, **e, g**), respectively. Molecular weight (**h-j**), charge at pH7 (**k-m**), and isoelectric point (**n-p**) of the poorly incorporated (Down, SV/raw (FC) < 1) and highly incorporated (Up, SV/raw (FC) > 1) proteins.

**Supplementary Data 5.**
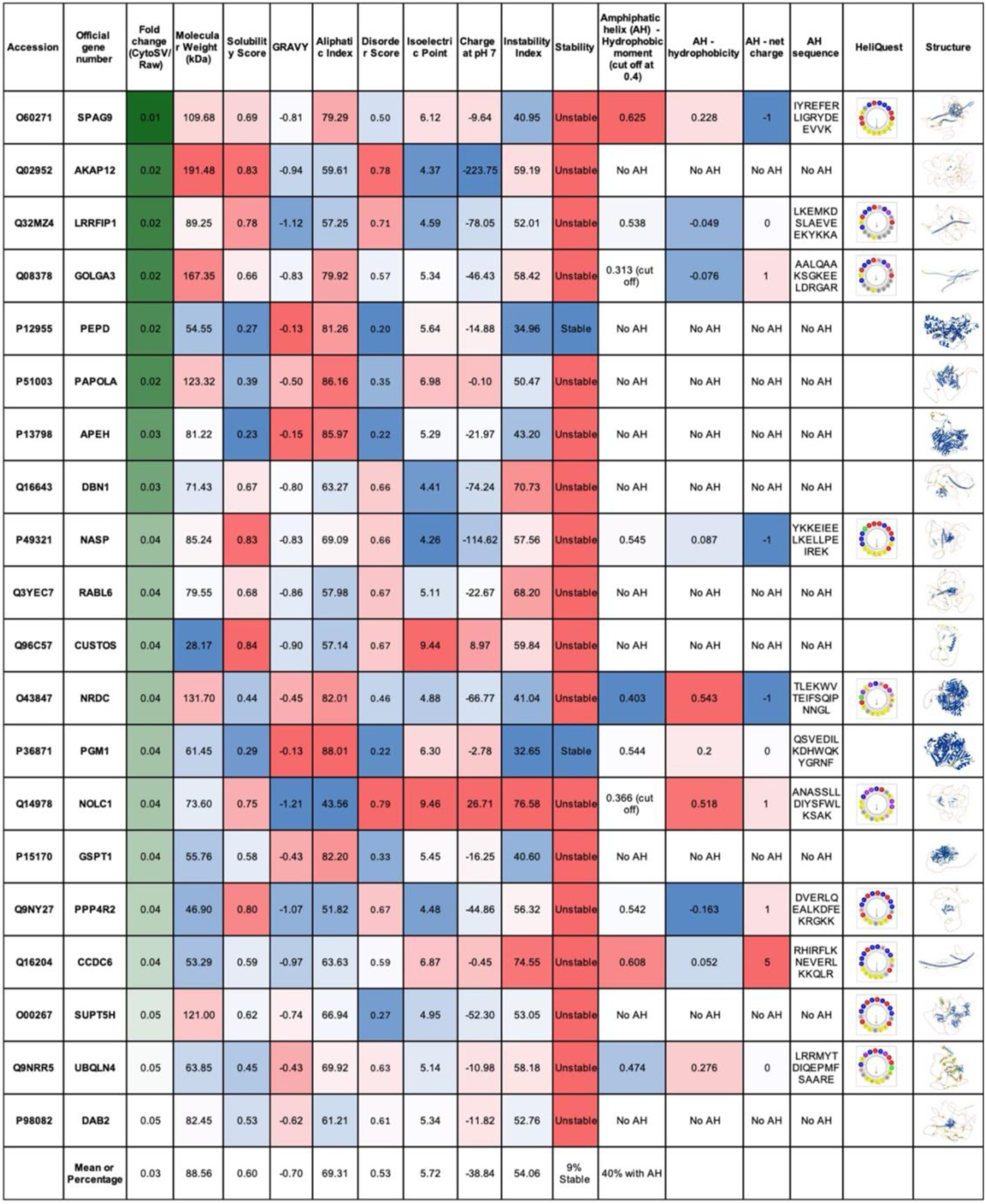
Characteristics of top 5% poorly loaded proteins in CytoSVs (Down in CytoSV). Color scale represents the gradient of the values within an individual column. Dark green to light green: Ranked by top most poorly incorporated proteins. Blue to red: low to high.

**Supplementary Data 6.**
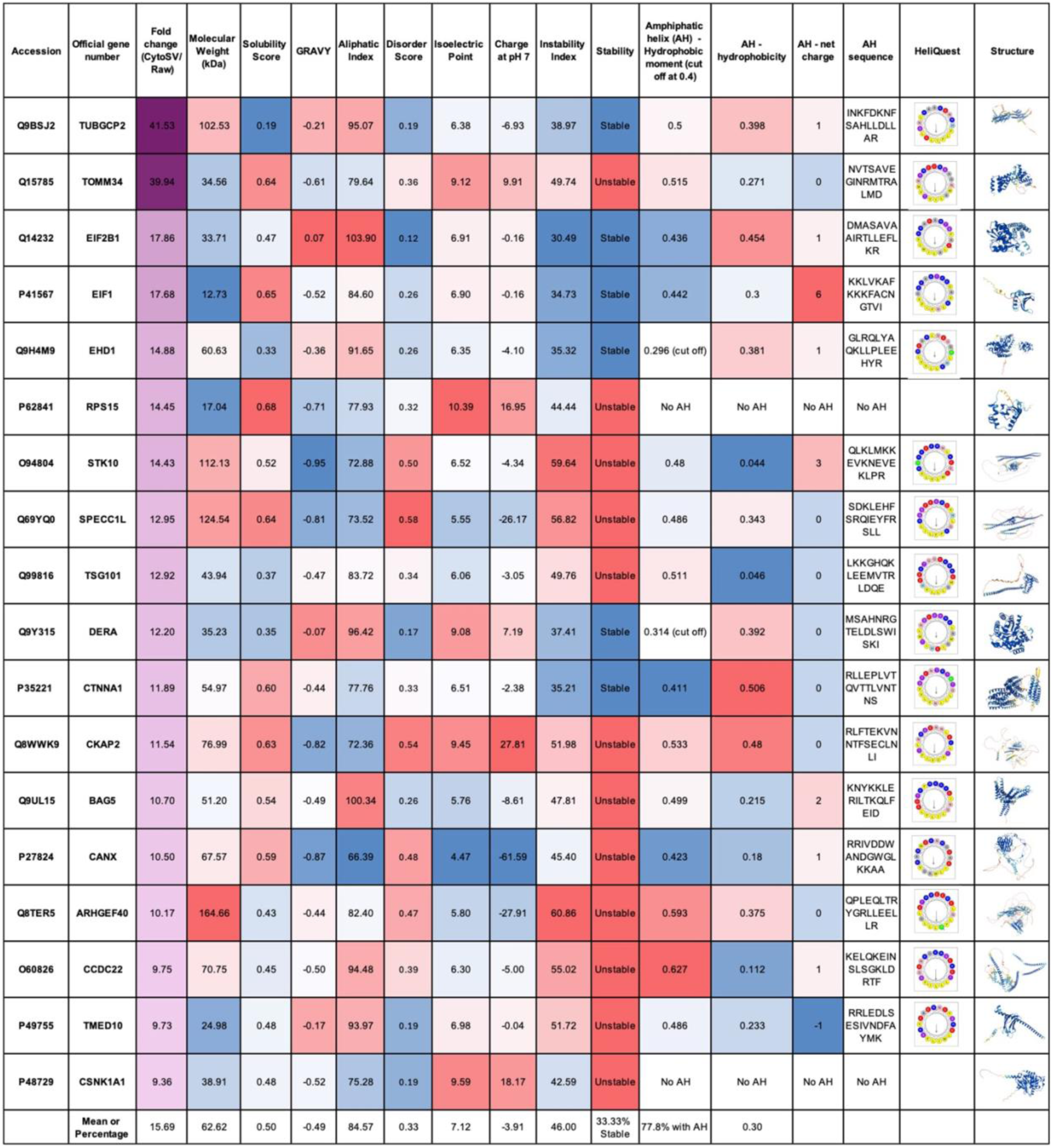
Characteristics of top 5% highly loaded proteins in CytoSVs (Up in CytoSV). Color scale represents the gradient of the values within an individual column. Dark purple to light purple: ranked by top most highly incorporated proteins. Blue to red: low to high.

**Supplementary Data 7.**
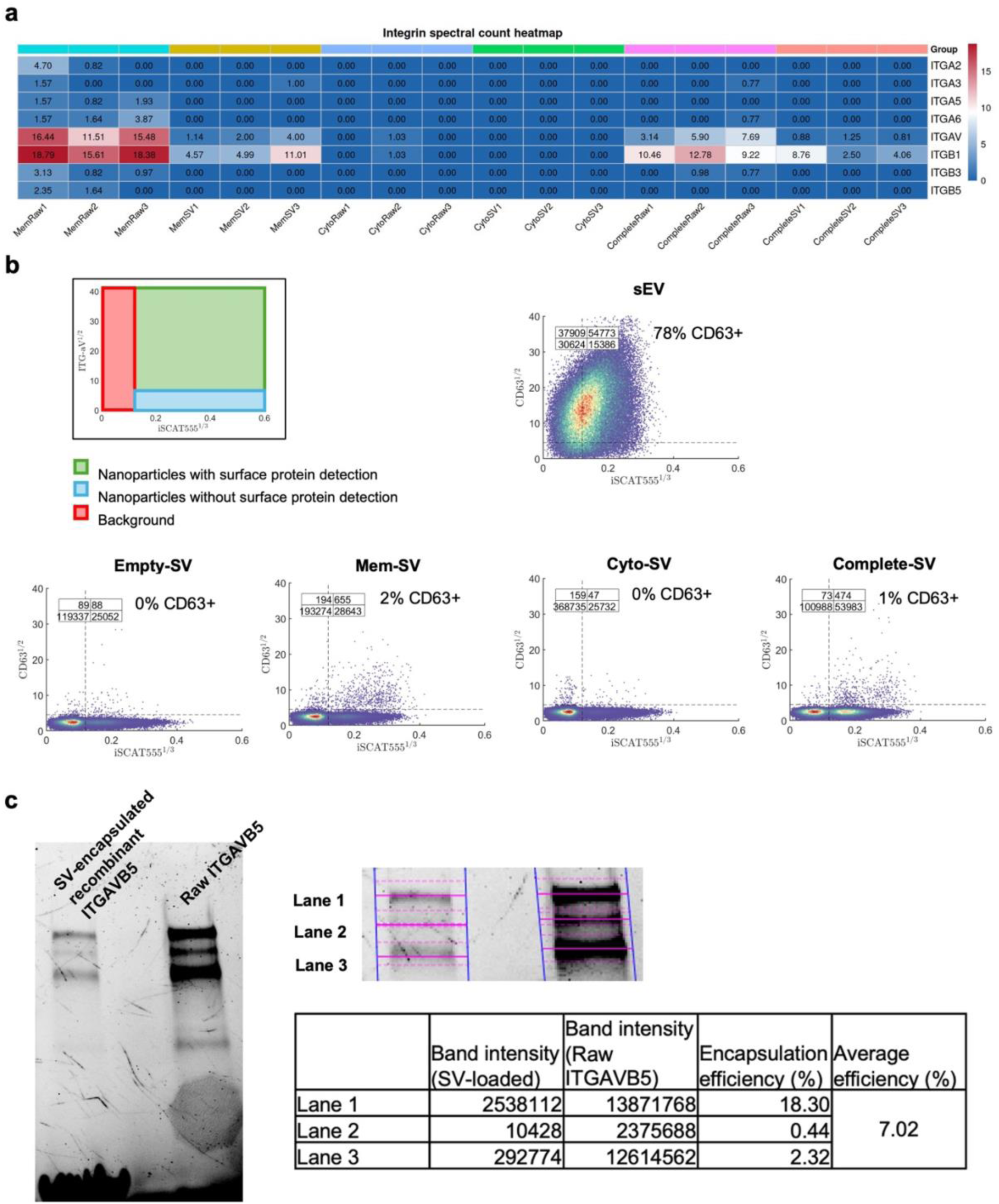
Supplementary data for individual protein quantification. **a**, Summary table of mass spectrometry spectral counts for integrins in the raw protein fractions across raw proteins and their corresponding SVs. **b**, Schematic illustrating how to interpret the SPFI (size photometry and fluorescence imaging^33^). CD63 detection on the surface of sEVs and SVs by SPFI. **c**, SDS-PAGE gel of recombinant ITG AVB5-loaded SVs and the raw ITG AVB5 equivalent to total loaded proteins. Estimation of encapsulation efficiency is performed using Image Lab.

**Supplementary Data 8.**
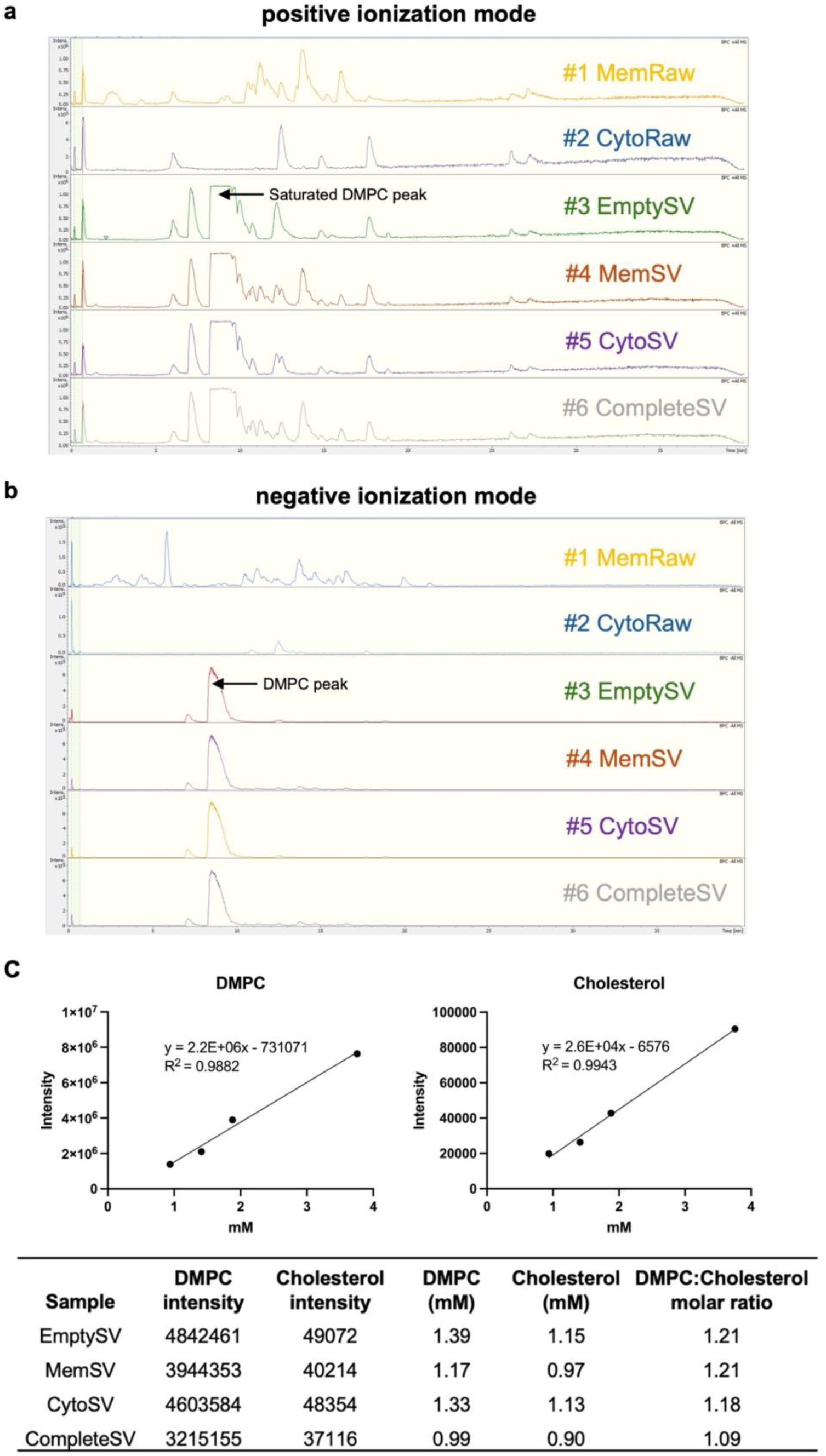
Extended lipidomics data. LC-MS chromatograms in (**a**) positive and (**b**) negative ionization modes. (**c**) Quantification of DMPC and cholesterol concentrations in SV samples using standard curves and dilution factors. Standard curves were generated from serial dilutions of synthetic DMPC and cholesterol with known concentrations.

**Supplementary Data 9.**
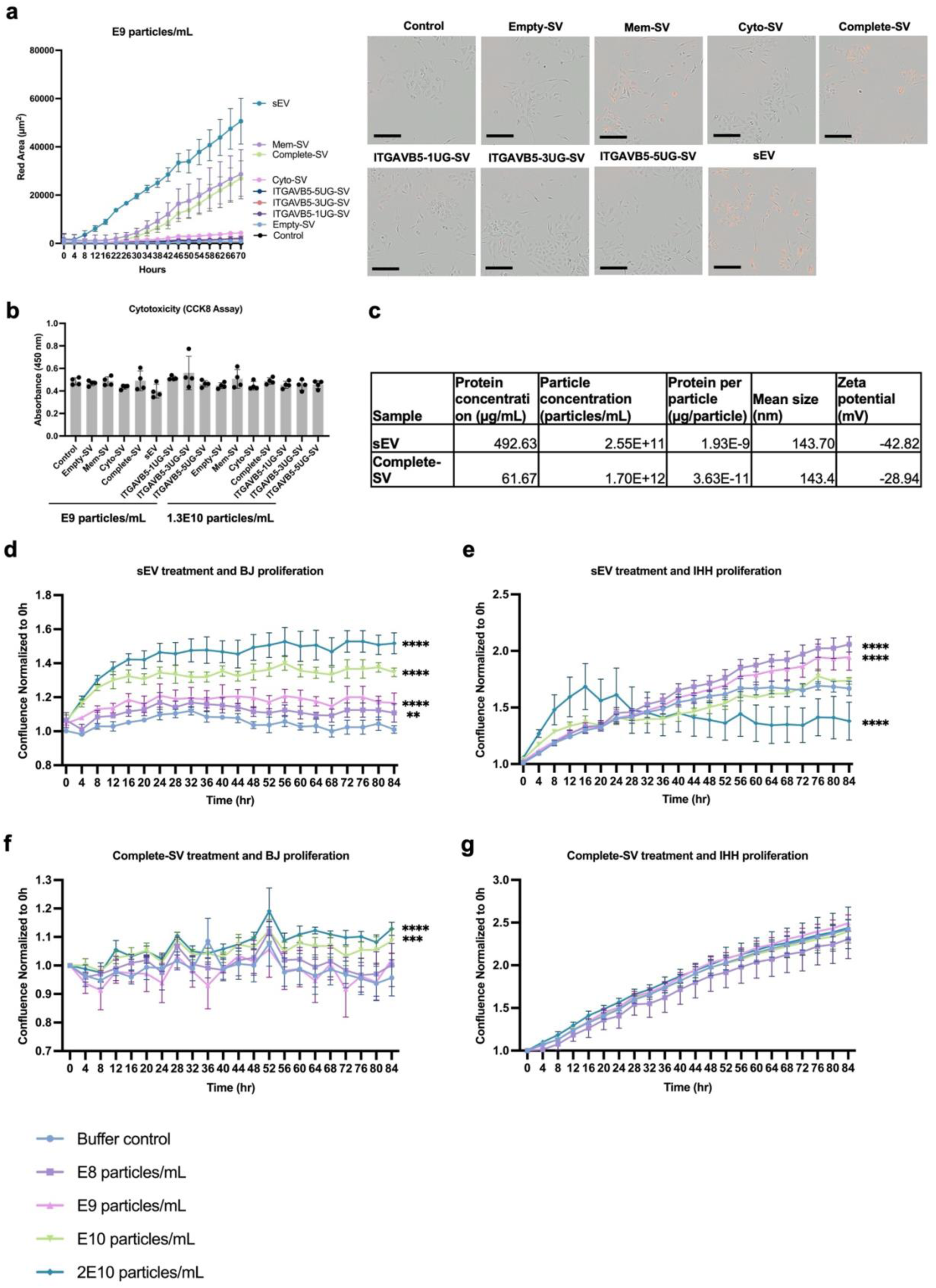
Effects of sEVs and Complete-SVs on BJ fibroblasts and IHH cells. **a**, IHH uptake of SVs and sEVs treated at E9 particles/mL (**b**) and uptake of sEVs treated at E10 particles/mL, and images at the endpoints were shown. Scale bars = 200 µm. Red fluorescence: sp-DiIC18 dye-labeled particles. **b**, Cytotoxicity assay of IHH by CCK8 assay post SV and sEV treatment. **c**, Comparison of the protein concentration, particle concentration, mean size, and zeta potential of MP41 sEVs and Complete-SVs. **d–e**, Effects of sEV treatment at increasing particle concentrations on the proliferation of BJ fibroblasts (**d**) and IHH cells (**e**). **f-g**, Effects of Complete-SV treatment at matched particle concentrations on the proliferation of BJ (**f**) and IHH (**g**) cells. Two-way ANOVA with multiple comparisons versus the control group (buffer control), with Tukey’s correction. Significance at the final time point (84 h) is shown. Mean ± SD. **, ***, and **** represent *P* < 0.01, < 0.001, and < 0.0001, respectively.

**Supplementary Data 10.**
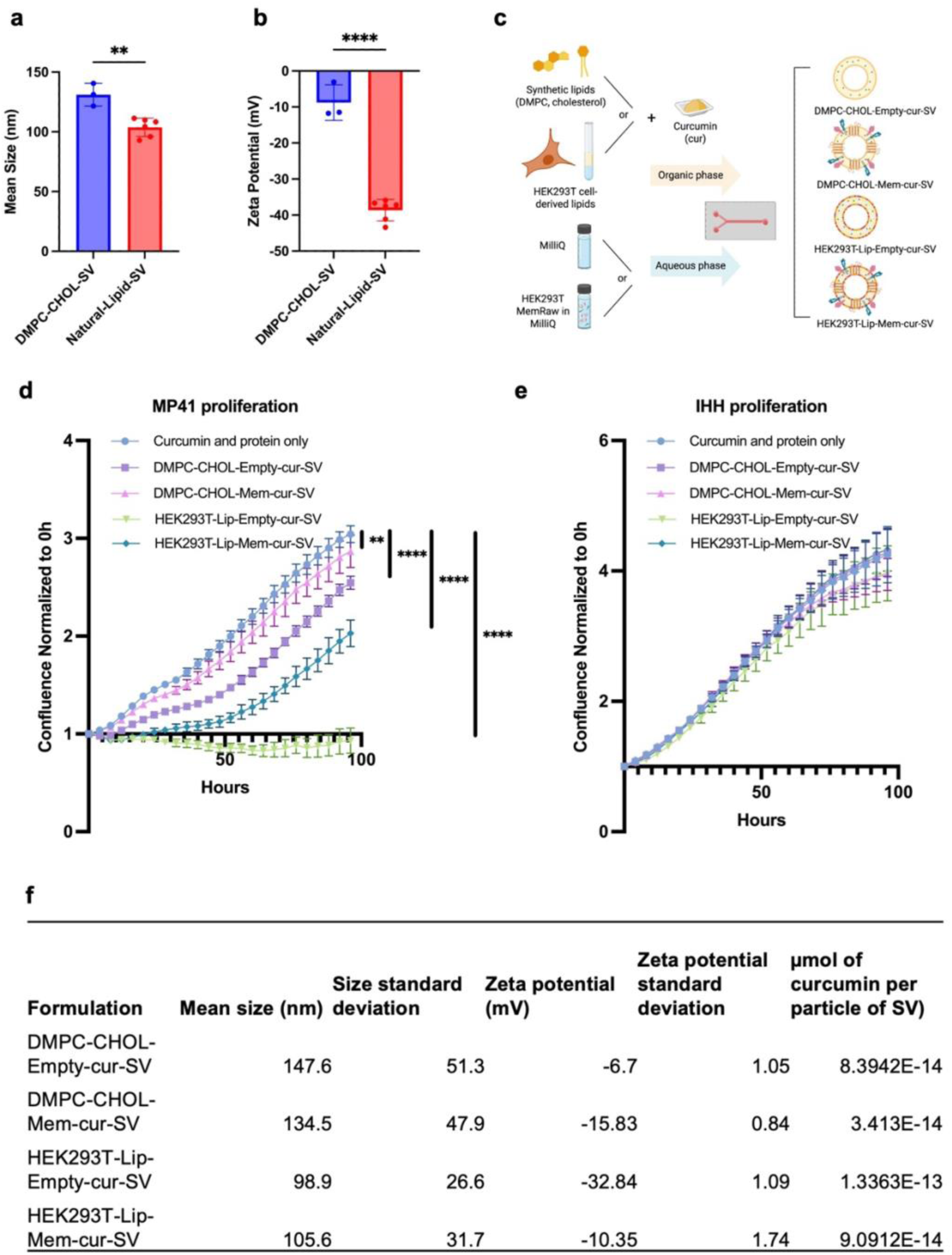
**Generalizability of SV fabrication using natural lipids and curcumin loading. a–b**, Comparison of particle size and zeta potential between SVs prepared from DMPC:CHOL (n = 3) and those prepared from natural lipid extracts (n = 6) of either cancer cells (MP41) or non-cancer cells (IHH, HEK293T), showing that natural-lipid SVs were smaller (**a**) and exhibited a more negative surface charge (**b**). **c**, Schematic of curcumin-loaded SV fabrication using either synthetic lipids (DMPC, cholesterol) or HEK293T-derived natural lipids and loaded with or without HEK293T protein sources. Real-time proliferation of MP41 (**d**) and IHH cells (**e**) treated with SVs, and the properties of which were summarized in the table (**f**), including particle size, zeta potential, and curcumin loading efficiency. (**d–e**) Two-way ANOVA with Tukey multiple comparisons versus the control group (curcumin and protein only), with Tukey’s correction. Significance at the final time point (96 h) is shown. Mean ± SD. **, and **** represent *P* < 0.01, 0.0001, respectively.

**Supplementary Data 11.**
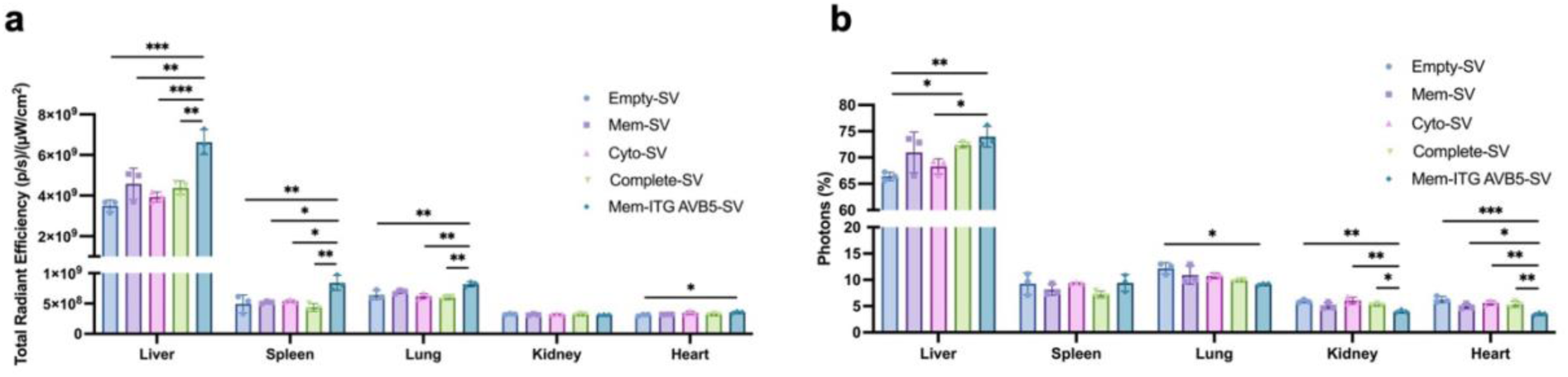
*Ex vivo* biodistribution data plotted in different metrics. Data for Fig. 6b plotted as total radiant efficiency (**a**) and percentage of total photons per organ over all measured organs (**b**). Mean ± SD. *, **, and *** represent *P* < 0.05, 0.01, 0.001, respectively.

**Supplementary Data 12.**
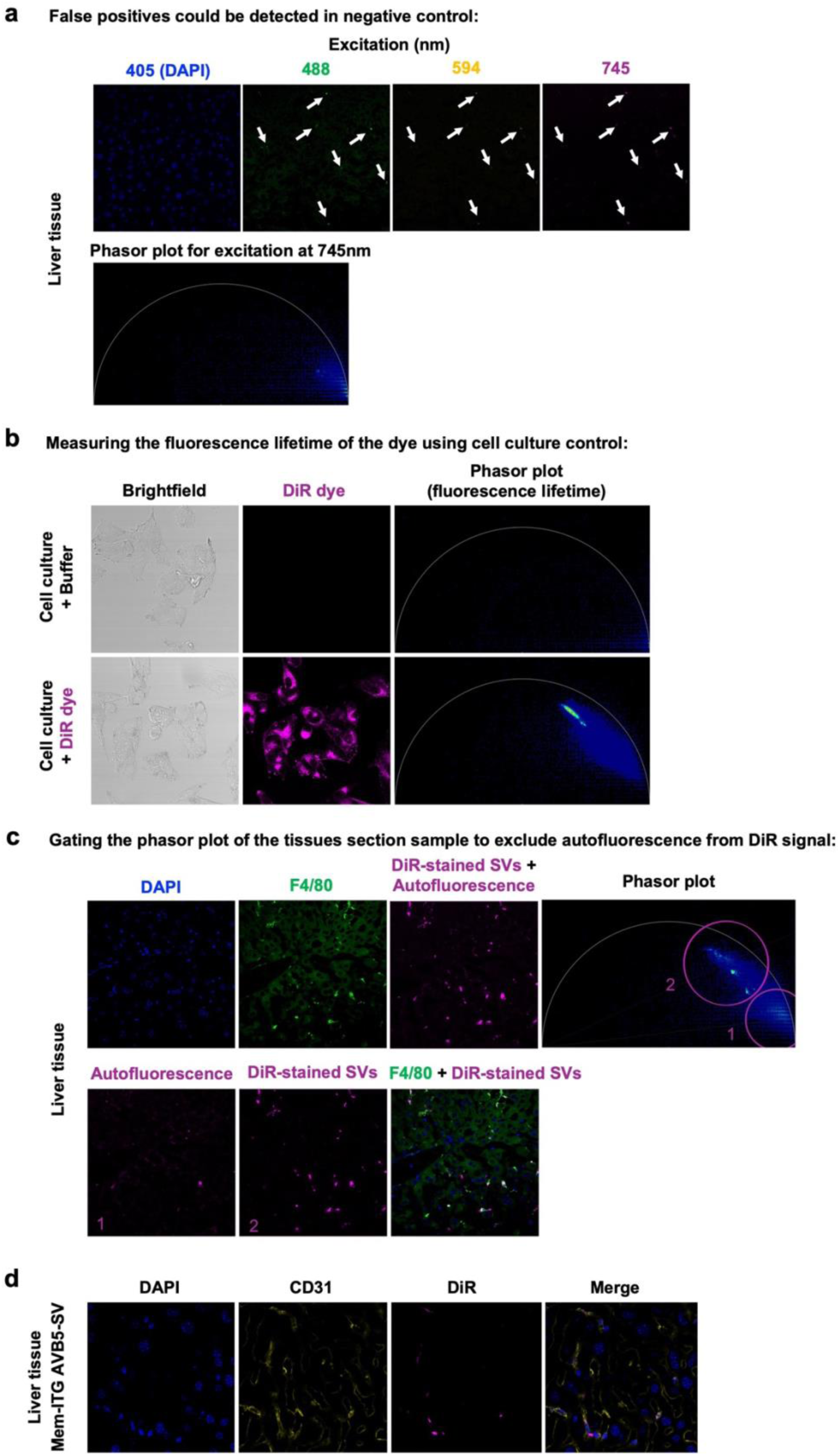
Removing autofluorescence by phasor plot in tissue imaging. Fluorescence imaging was performed using lasers with excitation wavelengths of 405 nm for DAPI, 488 nm for F4/80, 594 nm for CD31, and 745 nm for DiR. **a**, A liver tissue section from a negative control (mouse without SV or sEV injection) exhibited false-positive signals (autofluorescence) in the 488, 594, and 745 nm channels, showing overlapping patterns. A representative phasor plot of the fluorescence lifetime at 745 nm is displayed. To eliminate these false positives, which could interfere with co-localization analysis, the true fluorescence lifetime of DiR was determined using an autofluorescence-free cell culture (**b**). **c**, DiR signals (gate 2) from SVs (magenta) were distinguished from autofluorescence signals (gate 1) using a phasor plot, enabling accurate co-localization analysis with F4/80+ cells (green). **d**, Representative images depict the co-localization of Mem-ITG AVB5-SVs (magenta) with CD31+ cells (yellow) after autofluorescence removal.

